# Deep Mutational Scanning of structure-switching aptamers identifies both base requirement and base-pairing and uncovers regions required for either sensitivity or specificity

**DOI:** 10.64898/2026.09.06.749718

**Authors:** June H. Tan, Andrew G. Fraser

## Abstract

Structure-switching aptamers (SSAs) are short oligonucleotides that undergo a large conformational change on binding their specific target. This change can be used to read out target levels and SSA-based sensors have been developed for targets including drugs, metabolites, and toxins. Despite this potential as molecular sensors, we cannot predict how an SSA folds, how it binds its target, or the final SSA:target conformation and there are very few solved SSA structures. Here we use deep mutational scanning (DMS) to probe SSA structure and function, focusing on three published SSAs that detect cortisol. We show that DMS delivers rich data: it identifies which bases are required for activity but also maps base-pairing either in the presence or absence of target. We find all three SSAs have a similar structure when bound to their target despite having different predicted structures. To broaden our understanding of how SSAs can see cortisol, we selected many new cortisol-binding SSAs. Surprisingly, we identify diverse sequences that all have similar cores to the published cortisol SSAs — many diverse sequences thus converge on the same mode of target recognition. Finally, we use DMS to show we can change target specificity with a small number of mutations and map key specificity determinants. We conclude DMS can provide rich information on how SSAs work and that generating this for many SSAs could lead to greatly improved models for SSA mechanism.

## Introduction

Aptamers are short nucleic acid molecules that can each bind a specific target with high affinity and specificity^1,2^. Structure-switching aptamers (SSAs from here on) are a unique class of aptamer that undergoes a large conformational change on target binding. This conformational change can be used to drive a readout for target binding and SSAs thus have tremendous potential as molecular sensors^3^. Currently, there are SSAs that can recognise a wide range of targets including proteins, metabolites^4–6^, drugs^7^, toxins^8^, and even individual cations^9,10^. Each of these SSAs can be coupled to a molecular readout like fluorescence or a change in electrical conductance and used in simple devices. Alternatively, SSAs can be coupled to DNA barcode release allowing many different SSAs to be read in parallel using DNA sequencing in a recently described technique called smol-seq^11^.

Despite their great potential as molecular sensors, we understand very little about how SSAs work i.e. how they fold and how they recognise and bind their target. Every part of the functionality of an SSA is encoded in its sequence — the precise bases determine the structure of the SSA in the absence of the target, the affinity and specificity with which it recognises its target, and the conformation it adopts on target binding. Predicting SSA mechanism and structure from its sequence is a beguilingly simple problem — surely, it cannot be that hard to predict how a 30mer sequence will fold? Unfortunately this is very challenging indeed. First, even a short sequence like a 30mer can fold in many alternative ways that differ by only a very small amount of free energy — the energy landscape is much more rugged with many more local minima than protein folding landscapes. Second, there can be ‘kinetic traps’^12^ — this means that while one particular fold may be the most thermodynamically favourable, there may be no efficient route to achieve that structure and the SSA can be ‘trapped’ in one of these metastable intermediate folds. These are major problems that are computationally hard to solve and are only the first step: understanding how the SSA folds alone. There is then an additional tier to solve: how does the target dock to the SSA, which bases are required, and how does the SSA change conformation between free and target-bound conformations. Understanding how any known SSA works is thus extremely challenging and no algorithms routinely predict the correct structure of a SSA in complex with its target.

Recently AlphaFold has transformed protein structural biology — it is now possible to predict the structure of almost any protein to ∼atomic resolution within seconds given only the primary amino acid sequence^13^. However, it is not possible to do the same for structured nucleic acids like aptamers and riboswitches. AlphaFold was trained on rich training sets such as the solved structures of hundreds of thousands of proteins as well as deep evolutionary sequence data^13^. Such datasets are much more limited for aptamers including SSAs — there are very few examples where the structure of both the SSA alone and the SSA-target complex is known and solving structures is very slow. Is there an alternative way to generate rich experimental data to guide in silico predictions of SSA-target structures? Deep mutational scanning (DMS) has proved an excellent alternative way to identify interactions between amino acid residues and thus provide rich sets of spatial constraints that can be used to model protein structures^14,15^. However, DMS has been used surprisingly rarely in the aptamer field and here we wanted to test if it can shed light on SSA mechanism and structure as it has on protein structure and function.

Here, we use deep mutational scanning to examine the underlying mechanism of a set of three published SSAs that recognise cortisol but not highly related targets like progesterone or aldosterone^6^. We establish a pipeline that lets us examine the effect of every single mutation, every double mutation, and every deletion on SSA activity. We show that DMS can identify base-pairing interactions in SSA:target conformations and this indicates that all 3 cortisol SSAs recognise their target in a similar way. To expand this analysis, and test if there are other ways that SSAs can recognise cortisol, we carried out SELEX to identify many more SSAs that all recognise cortisol. We find many new SSAs that share highly similar cores to the published SSAs, suggesting that many diverse sequences converge on a similar mechanism of cortisol recognition. Finally, we show that DMS can be used to understand target specificity. We find that sensitivity to target and target specificity reside in different regions of the SSAs and that we can use DMS to discover the key bases required for specificity. We conclude that DMS of SSAs is a high-throughput technique that can be used to generate rich data on base-pairing interactions and conformational changes in known SSAs and that these can act as powerful constraints in building models of SSA mechanism.

### Outline of deep mutational scanning method for structure-switching aptamers

Yang *et al.* isolated a set of three SSAs that each recognise cortisol^6^ using SELEX from the same N30 library. Two of the three SSAs are predicted to form a three-way junction structure and have a highly similar core (CSS.1 and CSS.3) whereas the third (CSS.2) is predicted to have a distinct fundamental structure and core (Fig 1a). To better understand how these SSAs function, we wanted to use DMS to measure the impact of every possible single mutation, double mutation, and single deletion on the activity of these three SSAs.

**Figure 1:**
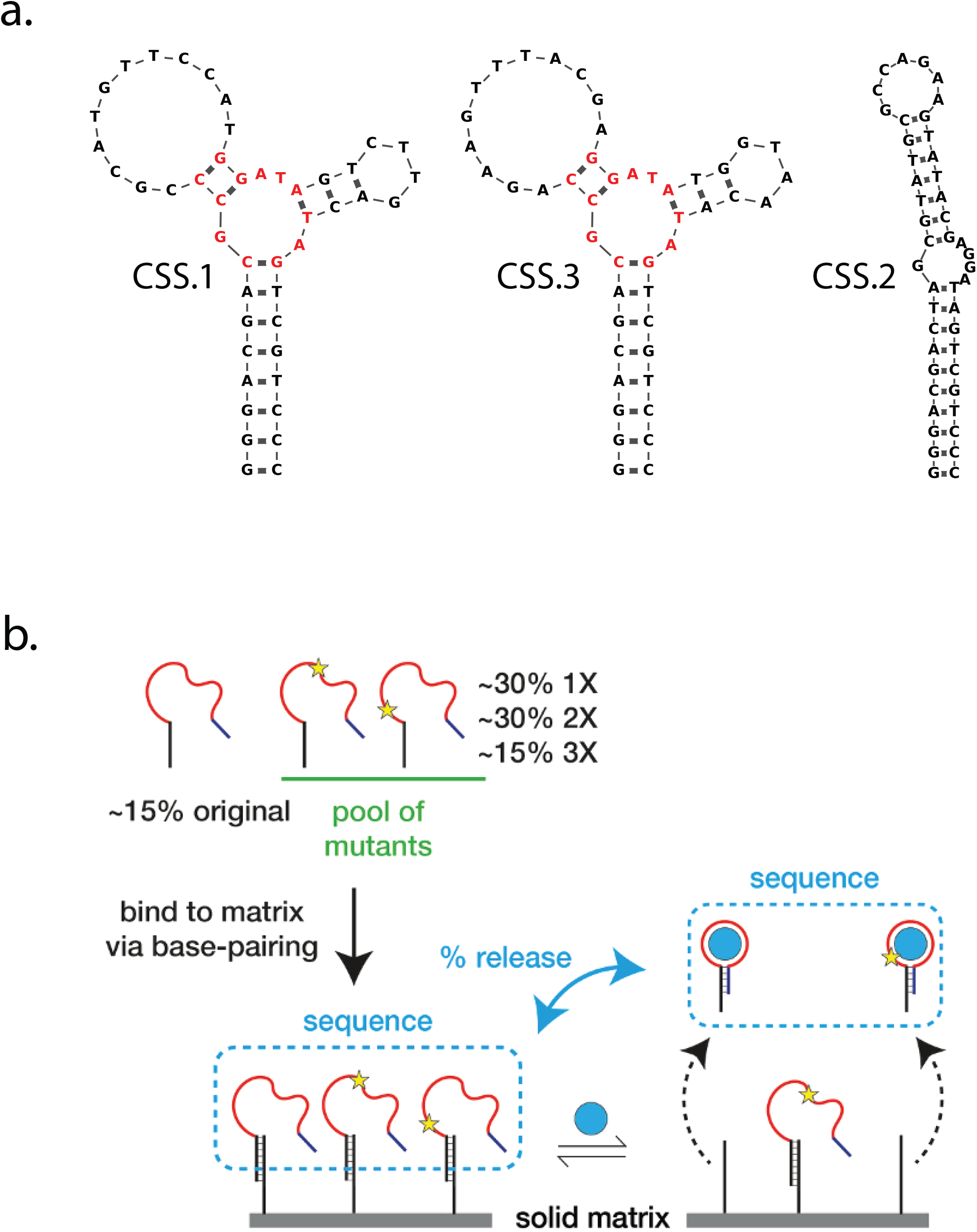
Deep mutational scanning (DMS) of cortisol SSAs. (a) Secondary structures of cortisol sensors (CSS.1, CSS.2, CSS.3) from Yang *et al.* (2017) predicted by Mfold. 8bp of the common stem region of the SSA is included in the structures. The identical core between CSS.1 and CSS.3 is highlighted. (b) Schematic of DMS of SSAs. A pool of SSAs with various mutations (star) is attached to a solid matrix via hybridisation with a short immobilised oligo which base pairs with the SSA oligo as shown. When a functional SSA binds its cognate ligand (blue circle), a conformation change induces the release of the SSA; those with inactivating mutations remain bound. Both released fraction and initial bound pool are sequenced to measure relative levels of release.

To generate the rich set of mutant SSAs we needed, we synthesised pools of oligos that contain mutations across the 30mer region that comprises the sequences required for target recognition. We do this using ‘doped synthesis’, that is, we use a 94:2:2:2 mixture of bases at each position during synthesis instead of the usual 100:0:0:0 ratio. For example if the first position is a C, 94% of the synthesised oligos will have a C at that position, but a random 2% will have an A, 2% will have a T, and 2% will have a G. Repeating this same process at each position of a known sequence results in a complex pool of mutant SSAs — for example, for a 30mer sequence ∼16% will have the correct original sequence, 30% will have a single mutation, 28% will have a double mutation and so on. In addition, ∼2% of our pool has a single deletion, due to imperfect coupling efficiency in standard phosphoramidite synthesis.

To screen this rich pool of mutants to identify which changes affect SSA activity, we first bind the entire pool onto a short immobilised oligo that base-pairs to the common stem (Fig 1b). We then expose the immobilised mutant pool to various doses of cortisol — active SSAs are able to bind the target and undergo conformational changes that drive release from the short oligo whereas mutants that are inactive cannot bind or release and remain immobilised. We capture the released active sequences and sequence them — this identifies every mutant that retains activity. Crucially, since we notice that mutations can affect the rate of SSA binding to the immobilised oligo and hence be over- or under-represented in the starting pool (Supp Fig 1), we also boil off all the SSAs that bound to the short immobilised oligo in a parallel sample — this measures how well each mutant sequence bound to short oligo. We can thus express the activity of every single sequence in the pool as a percentage release at each dose of target.

DMS identifies which mutations, or combinations of mutations affect SSA activity. The simplest result is the effect of single mutations: some can be mutated without any effect on activity, whereas others are absolutely required for activity. This is shown in Fig 2a-c for the three cortisol SSAs screened. In general, the data are simple: some bases can be changed without any effect, whereas others are absolutely required for activity (Supp Fig 2), but we note there are also a small number of bases which show strong sequence preference (Fig 2a-c), e.g. position 23 in CSS.2 can be C or T but not A or G. We also find the effect of single deletions to mostly reflect the effect of single base changes (Supp Fig 3). What this ‘single base’ view of the data does not reveal is any underlying mechanism — why are some bases absolutely required for activity? Are these bases required for ligand contact or are they required for structure? A partial answer comes from examining the effect of pairs of mutations on activity.

**Figure 2:**
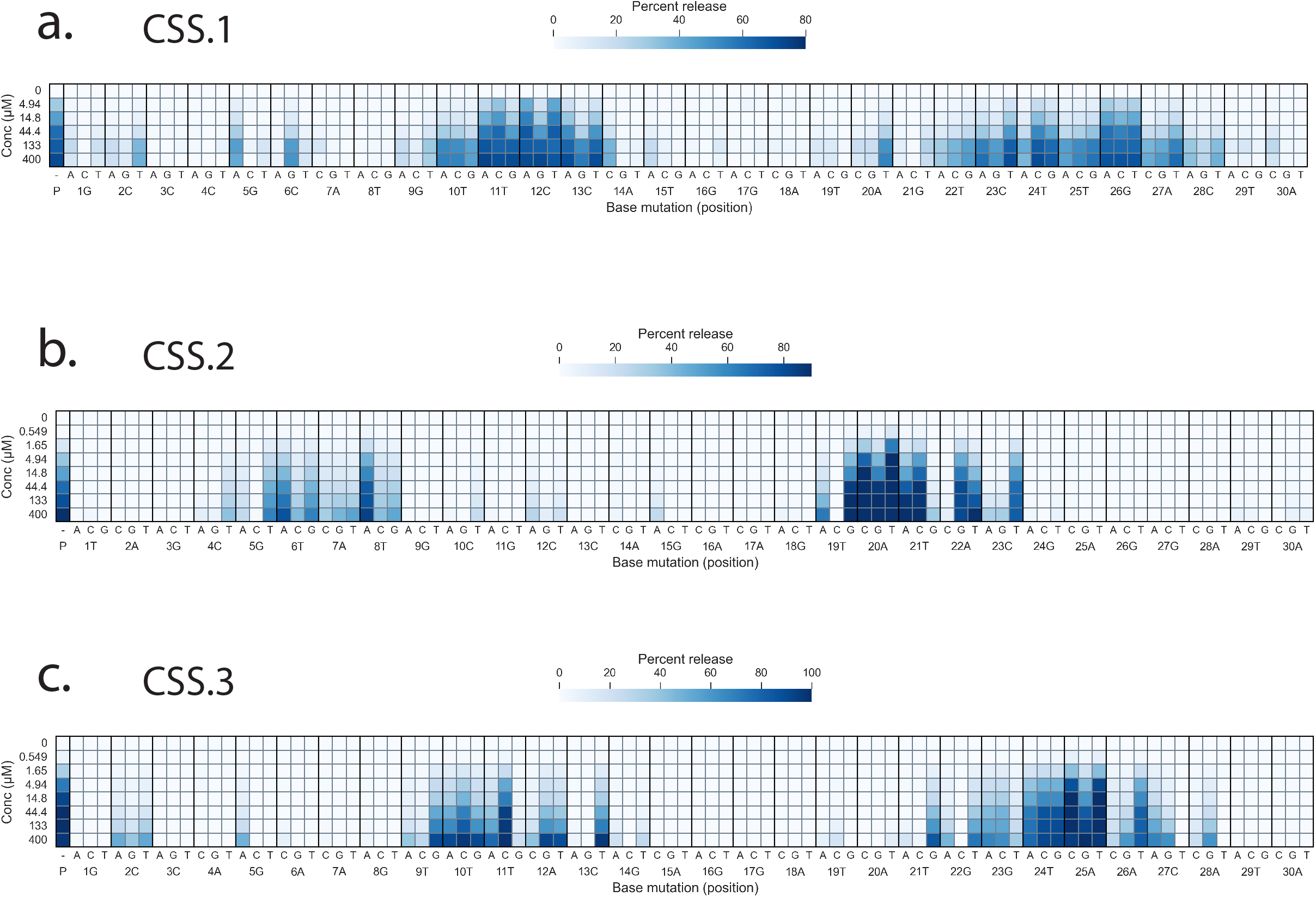
Effect of single mutations on cortisol dose response in CSS.1-3. The effects of each base substitution in the 30nt variable region of CSS.1 (a), CSS.2 (b) and CSS.3 (c) are shown. P denotes the response in the parental sequence and is plotted along with the effect of all possible single mutations. Columns show a dose response to cortisol in each of these sequences, with a darker shade of blue showing greater SSA activation. Percent release values were calculated from sequencing read counts of each sequence before and after ligand addition (details in Materials and Methods).

**Figure 3.**
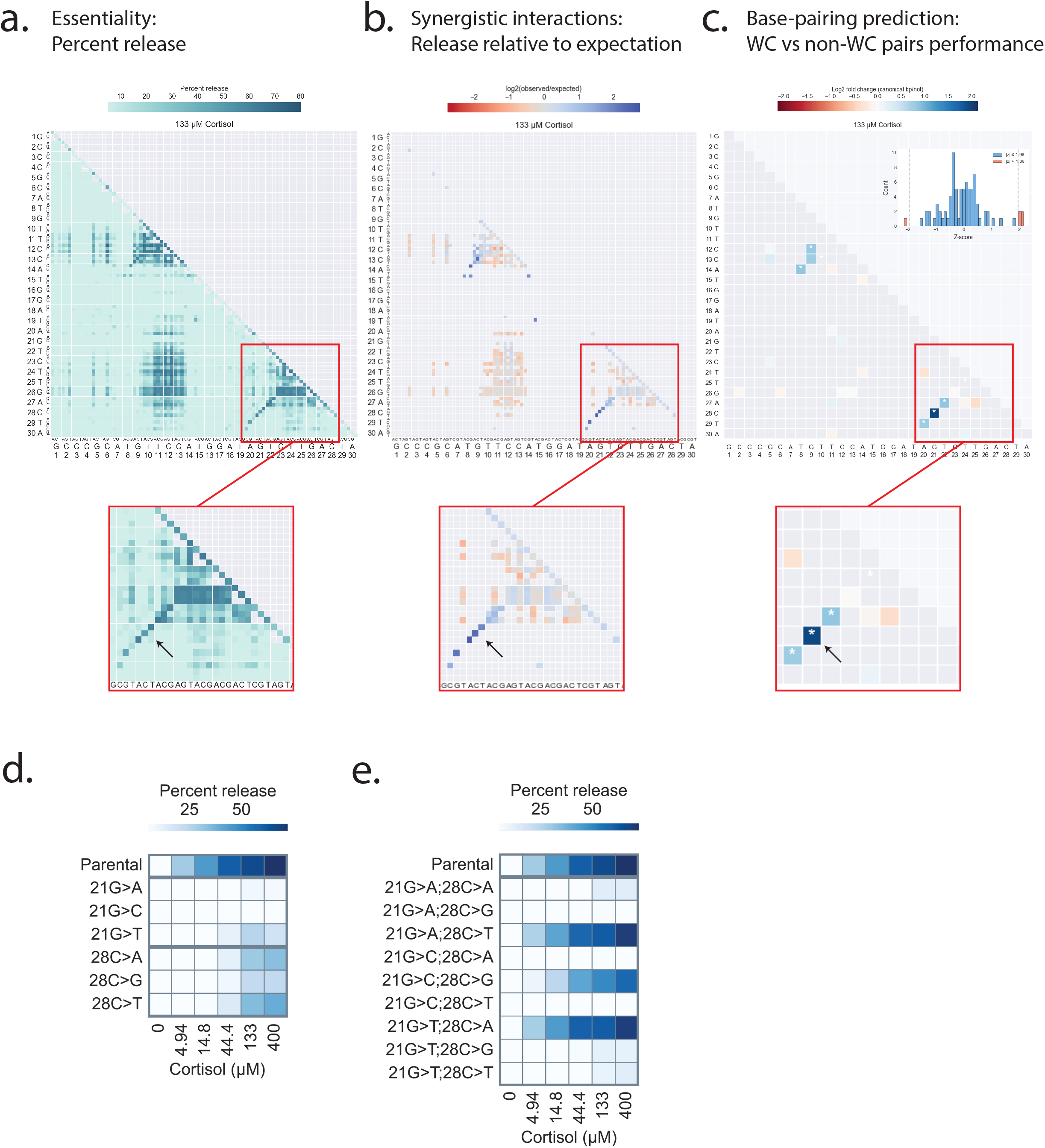
Mutation maps showing the effect of double mutations in CSS.1. (a) The effect of each double mutation is plotted, with a darker shade denoting higher SSA activation after incubation with 133 µM cortisol. Only double mutations with sufficient read counts are plotted. X-axis shows mutation 1 and y-axis shows mutation 2. Larger letters on the x-axis and y-axis represent the original base in the parent sequence while the smaller letters show the mutated base. (b) The activity of every double mutant is shown relative to expected activity given the effects of single mutations (multiplicative model). Only the effects of functional double mutant SSAs (>20% release) were calculated and plotted. Blue indicates the double mutant performs better than expected, while red indicates an effect worse than expected. (c) Comparison of double mutant activity at each position between standard Watson-Crick (WC) base pairs and non-WC base pairs. To test if bases at any two positions are likely to base pair, we averaged the interaction scores (from (b)) of double mutants that result in a WC base pair and compared it against the average interaction scores of the 6 other possible double base combinations. Positions where WC pairs perform better than other pairs are coloured in blue and an asterisk denotes a z-score at that position > 1.96. All two-position combinations were scored and used to calculate the z-scores but only positions where the original two bases are WC base pairs are coloured in. (d) Example of two essential bases in CSS.1; changing each base to any other results in greatly reduced activity. (e) Example of compensatory mutations between bases 21 and 28 where a second mutation that restores base-pairing (21A-28T, 21G-28C, 21T-28A) restores activity in the SSA. These two bases are highlighted by the black arrows in the double mutation maps in (a-c).

Fig 3 shows maps of how every possible pair of mutations affects activity. In raw form (Fig 3a), this simply shows the activity of each possible double mutant (given as percentage release), but this is not corrected for the effects of each single mutations: a combination of two single mutations that each has no effect on activity would be expected to have a very different activity than a combination of two mutations that each are essential for activity. A more useful view of the same data is in Fig 3b — instead of just plotting “percent release”, we ask: how different is the activity of the double mutant from what would be expected given the effects of each of the single mutations alone? We use a simple multiplicative model to predict expected double mutant phenotype i.e. activity of XY double mutant is expected to be activity of X single mutant * activity of Y single mutant — in Fig 3b, double mutations that are more active than would be expected from the individual effects of the single mutations are shown in blue, and double mutations that are less active are in red.

These maps show functional interactions between bases and yields clear information about canonical Watson-Crick base-pairing in the SSA in a similar way that phylogenetic data can reveal base-pairing in riboswitches^16^. For example, in CSS.1, position 21 is a G and position 28 is a C — both of these are essential and any mutations give an inactive SSA (Fig 3d). Since mutation of either base results in an inactive sensor, we expect all possible double mutations at 21 and 28 to be inactive. However, all double mutations that preserve base-pairing between 21 and 28 (i.e. 21A;28T, 21C;28G and 21T;28A) are all active (Fig 3e) — we thus conclude that there is a base-pair between 21G and 28C in the parental SSA that is required for function. Repeating this analysis for all possible base-base interactions identifies all Watson-Crick base-pairs in the SSA-target conformation (Fig 3c). We note that this approach cannot identify unusual base-base interactions that are often seen in aptamers like G-quadruplexes^17^ or triplexes and it will also fail to detect instances where bases pair but where there is some additional sequence requirement for one or other base. Nonetheless, these data show how DMS can give rapid insights into the canonical base-pairing that underlie SSA structures and this can be extremely useful in validating or excluding specific models.

CSS.1 and CSS.3 were previously proposed to have similar structures with a very similar core sequence (Fig 1a) and Mfold predictions of secondary structure are consistent with these published models — our DMS data largely agree with these proposed structures (Fig 4a-b). CSS.2, however, was proposed to have a very different structure and Mfold also predicts a very different base-pairing pattern to CSS.1 and CSS.3. However, our DMS data look very different to the proposed CSS.2 structure (see Fig 4a-b): we find base-pairing data in regions not predicted to be paired, and find no supporting base-pairing data in regions predicted to be stems. Remarkably, our DMS data suggest that the target-bound conformation for CSS.2 has the same conserved 3-way junction core as CSS.1 and CSS.3, but in a rotated variant where the universal stem takes the place of a hairpin loop and another hairpin takes the place of the stem (Fig 5a-b). When we align CSS.2 and CSS.3 around this 3-way junction elements, we also see 100% sequence identity among all essential bases in one of the loop regions extending from this 100% conserved core (Fig 5c), further supporting this alternative structure.

**Figure 4.**
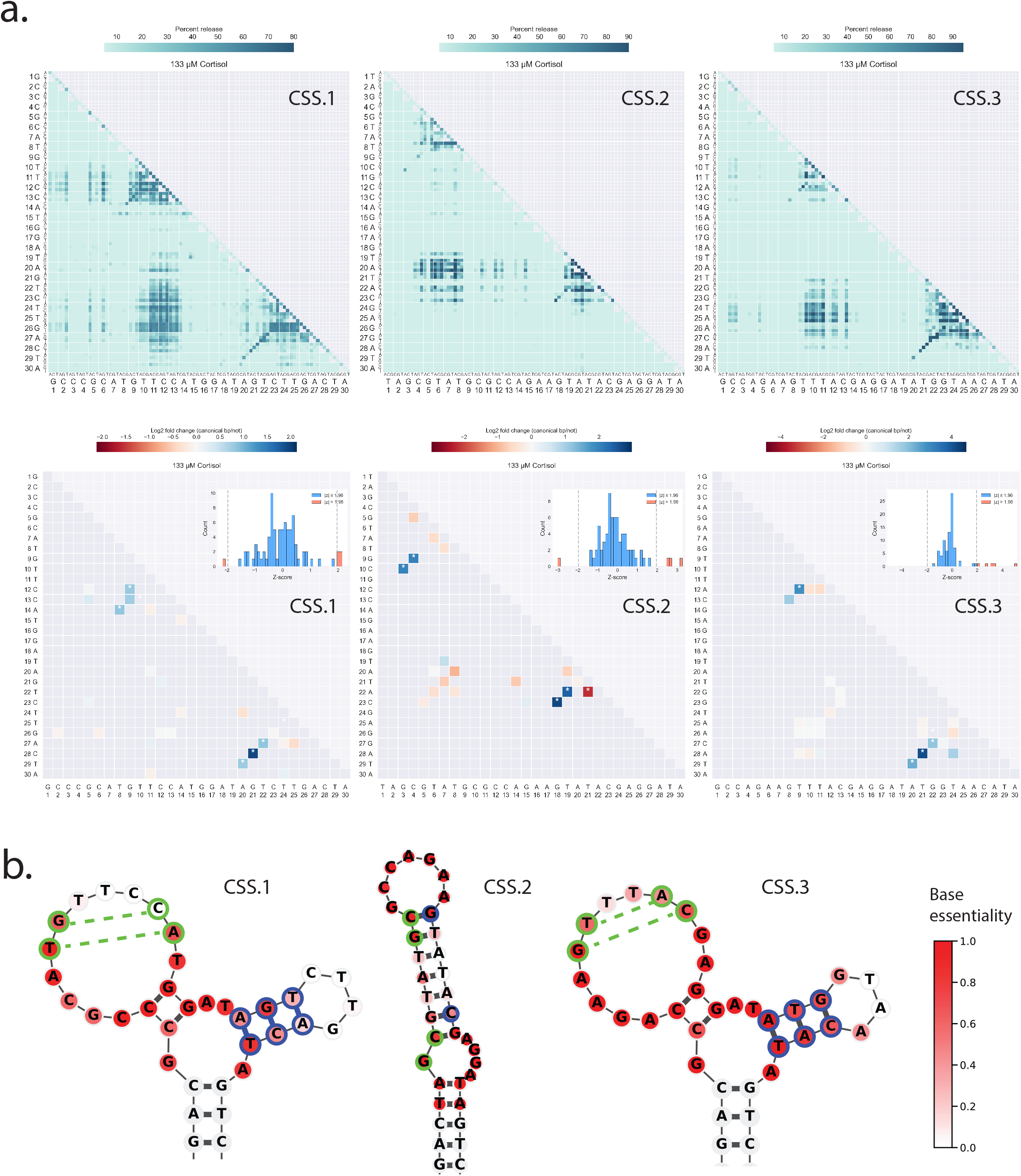
Double mutation profiles of CSS.1-3. (a) Double mutation maps of CSS.1, CSS.2 and CSS.3 at 133 µM cortisol are plotted and illustrated in the same way as described in Figure 3. (b) Single base essentiality and base-pairing predictions from DMS for CSS.1-3. Each base in the variable 30nt region of the SSAs are coloured based on the averaged response of single mutants at that specific position. Base essentiality is calculated as the inverse of percent release relative to parental. Red indicates an essential base where changing that base to any other results in greatly reduced SSA release in cortisol, while white indicates a non-essential base where changing the base results in minimal difference in SSA release relative to parental. Only 3bp of the common stem are shown here. Bases outlined in green or blue show sets of base-pairs inferred from DMS data.

**Figure 5.**
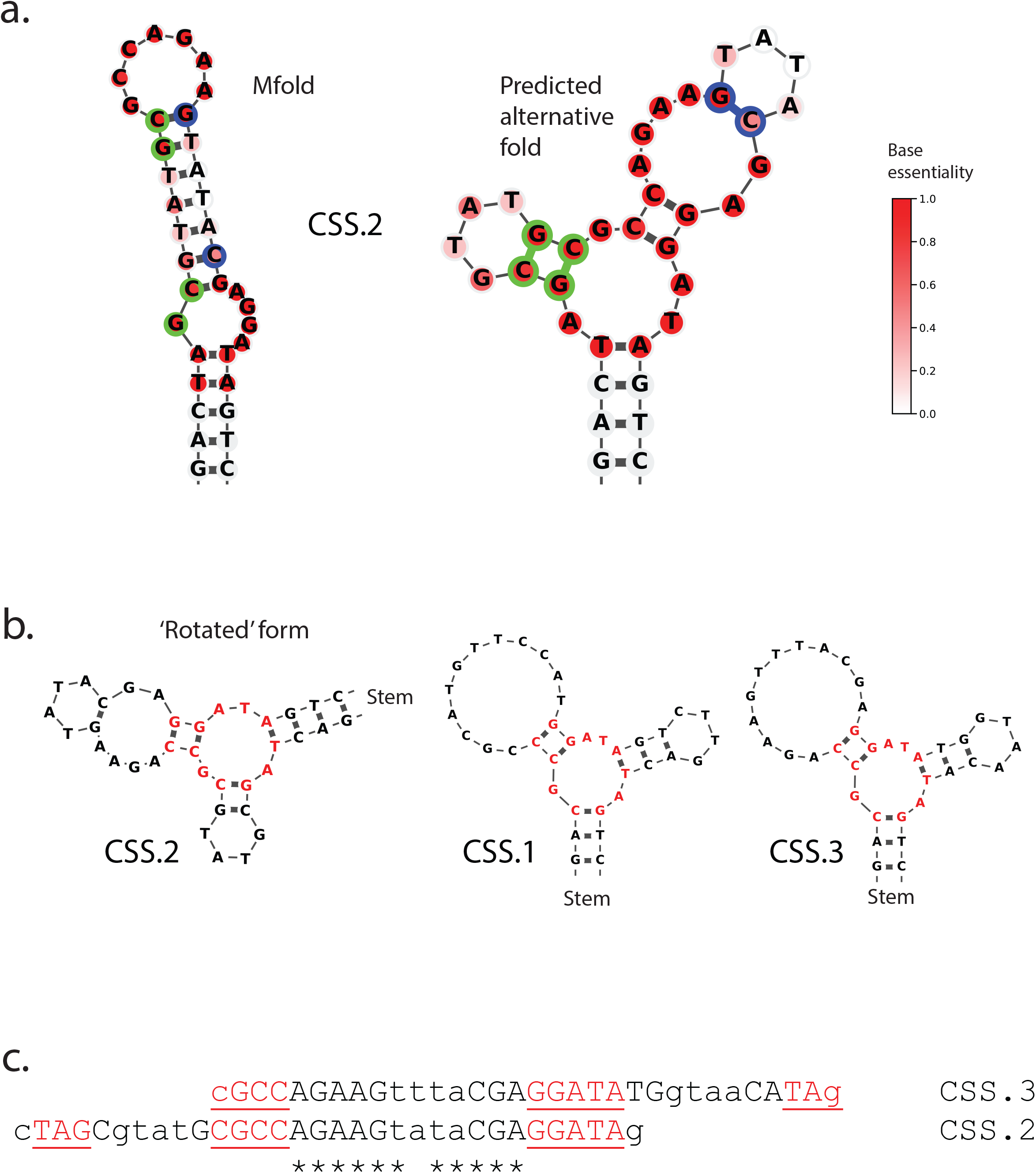
Possible alternative fold of CSS.2 inferred from DMS. (a) The structure prediction of CSS.2 from Mfold is illustrated beside an alternative predicted fold of CSS.2 based on DMS data. Only 3bp of the common stem region is shown here and each base is coloured based on essentiality from DMS data where darker red show more essential bases. Bases outlined in green or blue show sets of base-pairs inferred from DMS data. The alternative structure is predicted based on inclusion of DMS-predicted base-pairs and the common core seen in CSS.1 and CSS.3, and results in the less essential bases now being placed in hairpin loops rather than in the middle of the stem region in the Mfold-predicted structure. (b) The alternative predicted fold of CSS.2 contains the same common core as seen in CSS.1 and CSS.3 but in a rotated version where the common stem takes the place of one of the hairpin loops and another loop takes the place of the common stem. Identical bases at the 3-way junction core are highlighted in red. (c) Sequence alignment of CSS.2 and CSS.3 when aligned by core motifs. The two sequences are aligned by their shared motifs that is highlighted in red here and on their corresponding structures in (b). Bases in lowercase are bases in the universal stem bases or bases found to be non-essential from DMS. * denotes identical bases in one of the aligned stem-loop regions.

We conclude that deep mutational scanning is a rapid way to identify the key bases required for SSA structure and to identify many of the basepairs that are present in the SSA-target conformation.

### Identification of many new cortisol SSAs using SELEX

The SSAs CSS.1-CSS.3 were all identified using SELEX and all appear to bind cortisol with the same 3-way junction. Are there other possible SSA structures capable of responding to cortisol? During the CSS.1-CSS.3 selections, only a few SSAs were sequenced after many rounds of selections — there might thus be many different SSAs capable of detecting cortisol that were not identified in this process. To more comprehensively identify SSAs that respond to cortisol in a N30 library, we again used SELEX. We used an essentially identical N30 library as had been used by Yang *et al.*^6^ in their selection of CSS.1-CSS.3, but instead of only sequencing a limited number of identified hits after 15 rounds, we sequenced 20M reads from rounds 5-10 to identify as fully as possible the SSAs that responded to the target. We saw little enrichment of any sequences at round 5, but rounds 6-10 showed increasing enrichment of many sequences (Supp Fig 4). This yields many enriched sequences that are potential SSAs — each has to be retested and validated.

We retested and validated the potential new cortisol SSAs in two ways. The first was to re-order and retest individual sequences to confirm that they respond to cortisol as suggested by their enrichment during SELEX. The second was to carry out a replicate SELEX experiment and determine whether we see the same families of sequences being enriched in this independent SELEX.

We re-tested the 5000 most enriched sequences after 10 rounds of selection in a single pool to confirm that they are functional sensors — this is an efficient way to re-test the most enriched sequences. We find that only ∼5% of all 5000 sequences showed a significant response to cortisol (greater than 20% release at 133 µM cortisol). Most of the remaining ∼95% only responded very weakly to cortisol — while this weak response may explain their enrichment through 10 rounds of SELEX, it is not sufficiently strong to make them useable sensors.

We examined the ∼5% of SSAs that show >20% release at 133 µM cortisol to see if there were common features of these sequences. We first compared them at the primary sequence level — this immediately identifies near-identical SSAs that likely arose by PCR errors during repeated amplification of an initial parent sequence and we find some examples of these (Supp Fig 5a). However most of the identified sequences share no strong primary sequence similarity and are ‘orphans’ when clustered by sequence. To gain more insight into whether these shared higher level similarity, we compared the secondary structures predicted by Mfold, focusing on ‘open’ structural elements such as 3-way junctions and internal loops. This revealed clusters of sequences that share the same core – many of these were not previously clustered together by sequence similarity (Supp Fig 5b). 127 of the 277 SSAs that show activity at this threshold are predicted to form 3-way junctions and 79 of these 127 3-way junction SSAs fall into 6 clusters of sequences (Supp Fig 5b). We then filtered out similar hits that were likely derived from PCR errors and thus not completely independently selected for. Examples of the resulting diverse members of each of these 6 top cluster are shown in Fig 6a along with their unique core and dose response; Fig 6b additionally shows the response of individual representatives of each cluster to cortisol and a bile acid, tauroursodeoxycholic acid (TUDCA), that is structurally-related to cortisol to test both sensitivity and a measure of specificity. We note that members of each distinct cluster tend to have similar responses and specificities confirming that each cluster represents a set of sequences that function similarly.

**Figure 6.**
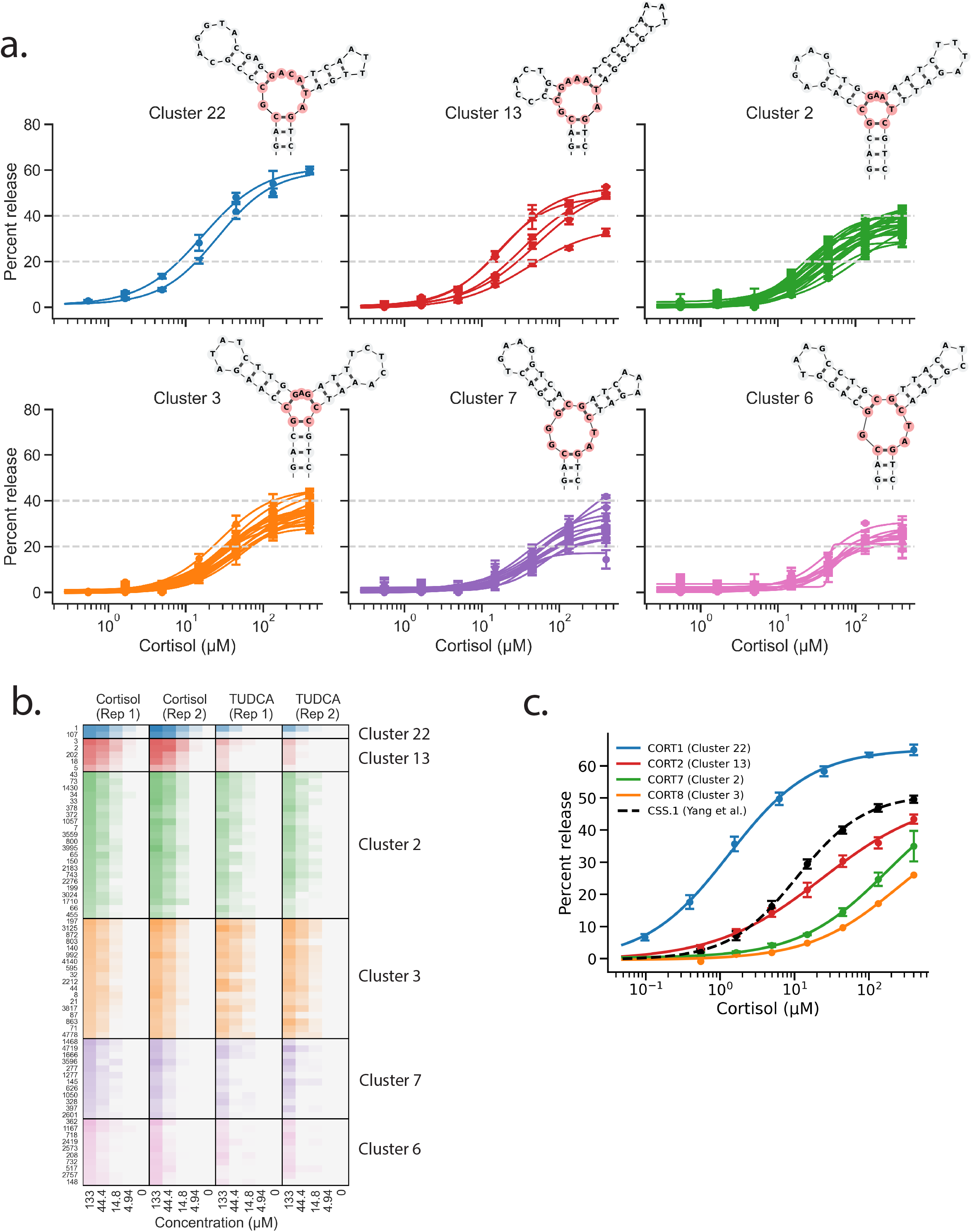
Top 6 performing clusters from SELEX. (a) Dose response curves of sequences in the top 6 SELEX clusters. All sequences belonging to these 6 clusters and among the initial top 5000 hits were first filtered to remove closely related sequences (Hamming distance of 2) and the activity of the filtered individual sequences were then plotted. Two replicates were carried out for these sequencing-based re-tests and error bars denote standard error. The Mfold-predicted structures of the top performing representatives in each cluster are illustrated. The junction core used to cluster and group these sequences are highlighted in red. (b) Dose response profiles in the presence of cortisol or TUDCA. Sequences from each cluster were first filtered as in (a) and their dose responses in either target ligand were then plotted for comparison. (c) Dose response curves from individual fluorescence-based assays. Representative sequences from the top 4 clusters were re-tested as individual sensors in a fluorescence-based assay. In this case the SSA is immobilised and hybridised to a short 6-FAM-attached oligo, which is released and read out on ligand binding (see Materials and Methods for details). The dotted line shows the dose response of the CSS.1 sensor in this assay and architecture for comparison.

Two clusters showed at least as strong responses to cortisol as the published CSS.1, CSS.2, and CSS.3 — clusters 13 and 22. These are the only two clusters with SSAs that give >40% release at 400 µM cortisol and some members show greater sensitivity than the published sensor (Fig 6c). Sequences in these two clusters comprise all of the top 22 hits among the top 5000 hits we re-tested and these two clusters have a core that is very similar to CSS.1-3. Both cluster 13 and cluster 22 share a 3-way junction core that is almost identical with the published CSS.1 3-way junction core, with the exception of one free base in the core (a T in CSS.1-3 that is a C in cluster 22 and an A in cluster 13 (Fig 7a). In total, we identified 7 diverse members of this class of SSA (<70% sequence similarity to any other) — 2 from cluster 22 and 5 from cluster 13.

**Figure 7.**
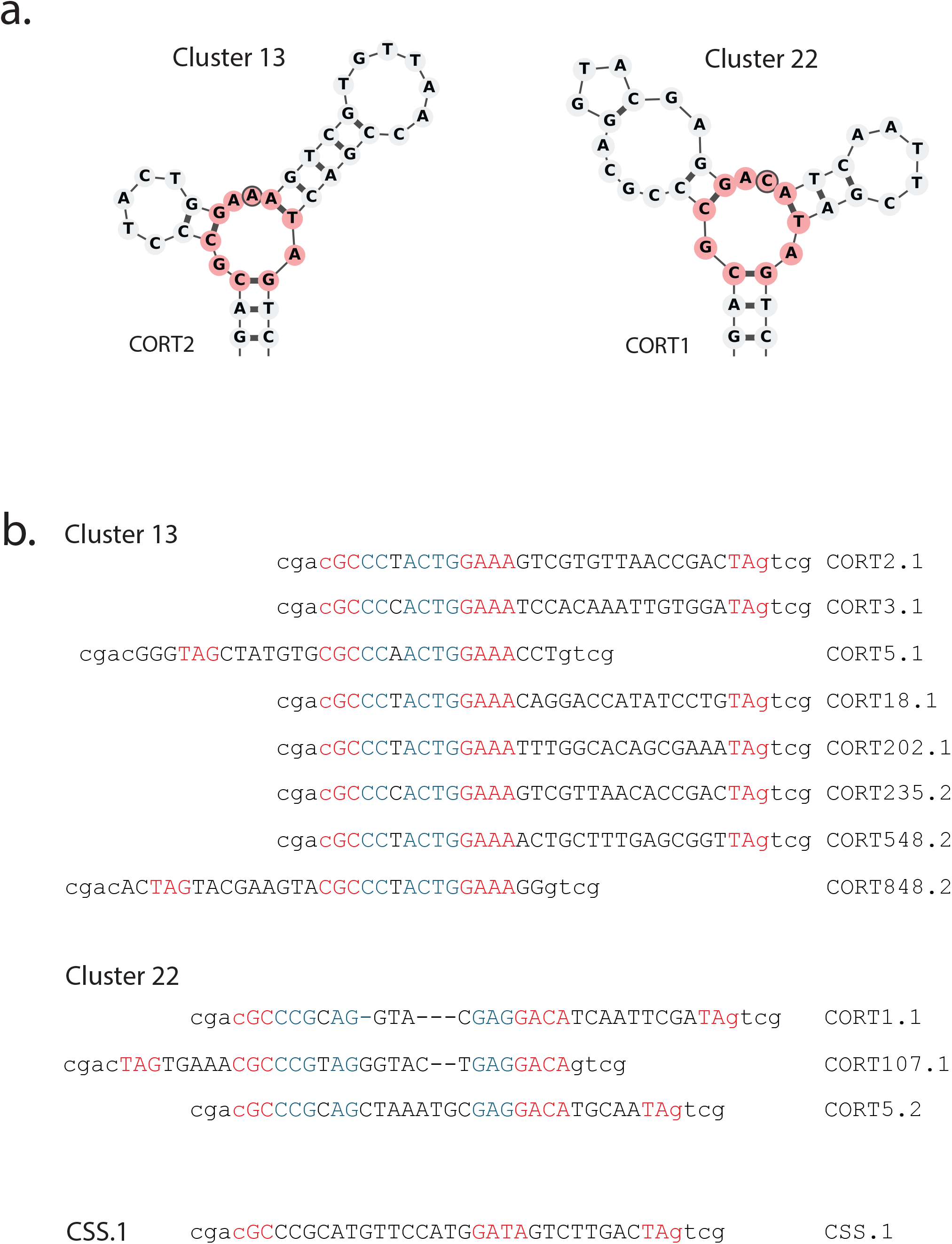
Two SELEX clusters show similarities to the known cortisol sensors. (a) Representative structures from Cluster 13 and Cluster 22 as predicted by Mfold or manual annotation based on DMS data. The common cores of sequences in these clusters are highlighted in red. The bases in the 3-way junction core that differs between these clusters and CSS.1 are outlined in black. (b) Sequences in Cluster 13 and Cluster 22 are obtained from two independent SELEX experiments and aligned based on core motifs. ‘.1’ at the end of the sequence name denotes a sequence obtained from the first SELEX, while a ‘.2’ is from the second SELEX experiment. The junction core bases used for cluster assignment are highlighted in red while stretches of consensus bases in a stem-loop region is highlighted in blue. Lowercase letters refer to bases in the universal stem region which is fixed among all sequences in the SELEX pool.

We next carried out a second independent SELEX experiment using cortisol as a target to determine how reproducible SELEX is and whether we find different sets of sensors. What we find is striking —this second independent SELEX yields very similar results and we identify new members of cluster 13 and 22 in the most enriched 1000 sequences. In total, after these 2 independent SELEX experiments we have 3 distinct members of cluster 22 and 8 members of cluster 13 and we focus on these two clusters for the remainder of this paper as they are the most robust and sensitive sets of SSAs that we isolated.

Comparing these new 11 cortisol SSAs to the published CSS.1-CSS.3 cortisol SSAs identifies both common and different features. First, the common region of these new 11 SSAs extends beyond the CSS1-like core, and includes an additional loop extending to one side of the junction (Fig 7a). Almost all the SSAs we identified in cluster 13 have near-identity across this short loop while we also see stretches of consensus bases among SSAs in cluster 22 (Fig 7b). This is intriguing — while the high conservation over this additional region in our new cortisol sensors suggests that it has some function, there is no conservation over this region between these new SSAs and CSS.1, or between the two new clusters we identified. Second, many quite different sequences can result in the same consensus core (Fig 8a) — in particular there is a stem loop structure in all the SSAs that extends between a conserved GA(A|C)A and a 100% identical TAG. That stem-loop can have different stem sequence composition, the loop has no obvious sequence constraints and can be of various lengths, and the stem can also contain bulges without any major consequence on function (examples in Fig 8a). More remarkably, just as we postulated for CSS.2 in Fig 5b, even the position of the universal stem can vary between members of this family of SSAs. As shown in Fig 8b and Fig 8c, we isolated rare examples of SSAs from both cluster 13 (Fig 8b) and cluster 22 (Fig 8c) where the universal stem and unconstrained stem-loop have switched positions — ultimately, the core is the same, the positions of stems is the same, but the identity of the stems has shuffled. This flexibility in both architecture and primary sequence in unconstrained regions explains why SSAs with little sequence identity can still use the same cores to bind their targets. We conclude that rich SELEX data can identify the functional cores of related SSAs.

**Figure 8.**
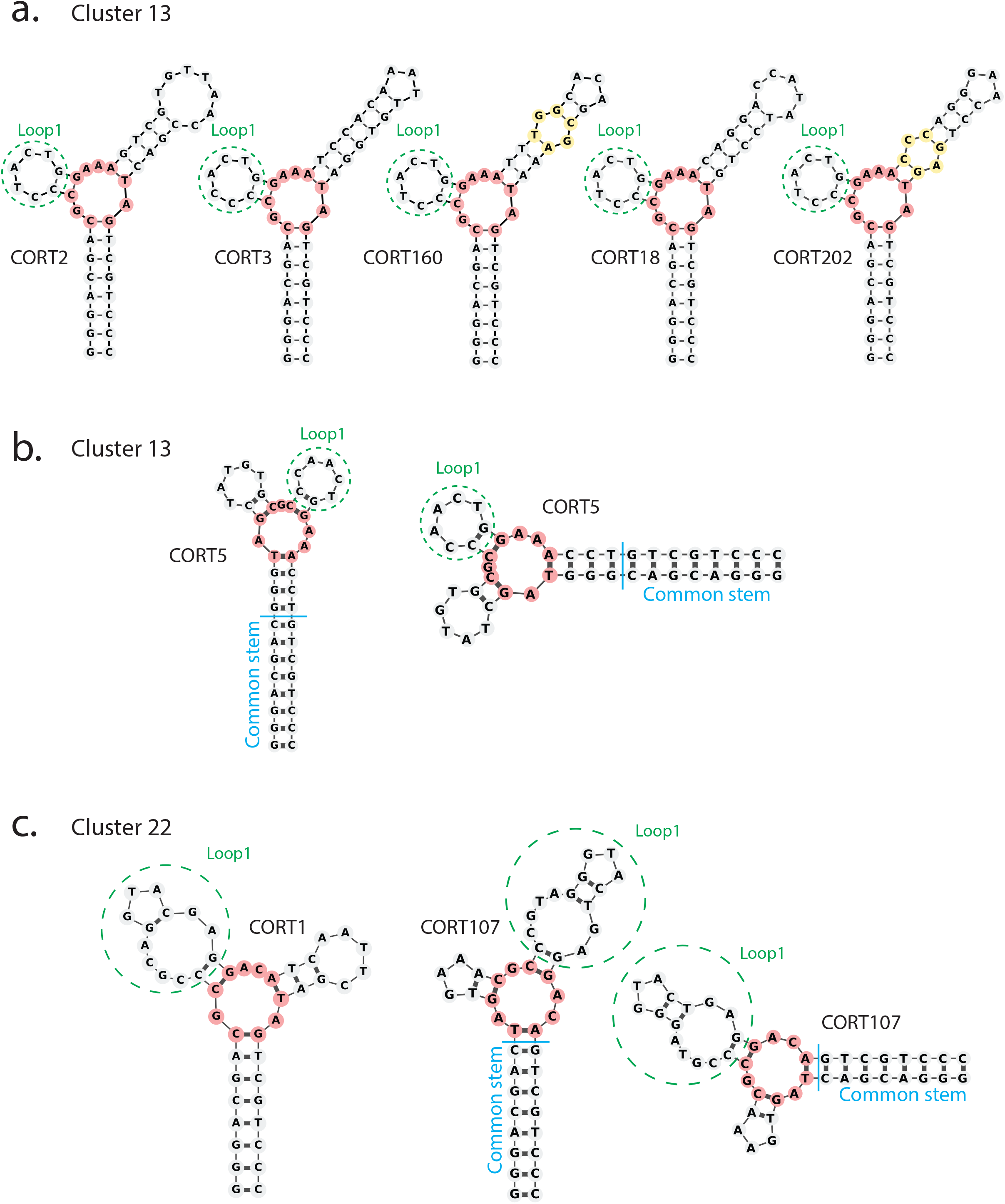
SSAs in clusters 13 and 22 share a highly similar stem-loop region among sequences in their cluster. (a) Sequences in SELEX cluster 13 share a strong consensus in one of the stem-loop regions circled in green (referred to as ‘Loop1’ from here on out), in contrast to the limited sequence similarity seen in the other stem-loop. The secondary structures predicted by Mfold are shown. The cluster core is highlighted in red while internal loops and bulges in stems are highlighted in yellow. (b-c) The position of the universal stem can vary between cluster members. CORT5 (b) shows an example of a rotated version of the junction core of cluster 13 where the common stem and loops have switched positions around the 3-way junction, but CORT5 still shares the same consensus in ‘Loop1’ as other cluster members in (a). CORT107 (c) shows an example of a rotated version of the core of cluster 22 where ‘Loop1’ has high sequence similarity with other cluster members (CORT1). The secondary structures of CORT1 and CORT107 shown here are based on a combination of Mfold predictions and manual structure refinement using DMS data.

### Deep mutational scanning and rich SELEX data identify the same consensus core

The rich data obtained from our SELEX selection of SSAs that respond to cortisol defines potential consensus sequence and structural elements. To identify these types of elements more comprehensively, we used DMS to examine the sequence requirements for activity of two SSAs in cluster 13. The results are strikingly similar to the SELEX data — the bases identified as being essential by deep mutational scanning are highly conserved between members of the SELEX cluster and the predicted base-pairing from Mfold is largely accurate (Fig 9). SELEX and DMS thus converge on the same insights into how a family of related SSAs works, which bases are required for function, and the likely base-pairings in the SSA. SELEX samples a far larger sequence space while DMS comprehensively covers a smaller sequence space around an initial parental anchor sequence.

**Figure 9.**
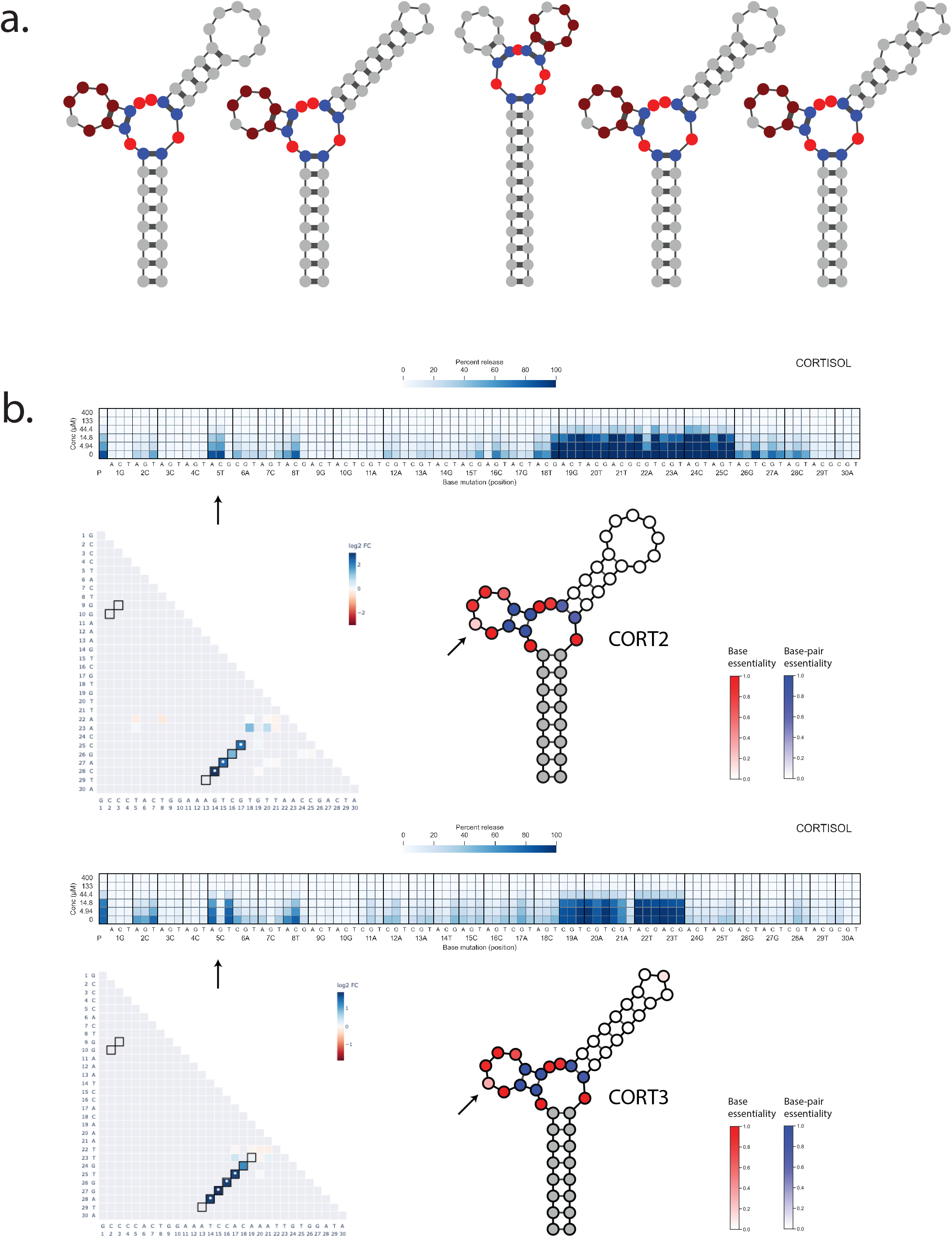
DMS and SELEX data identify the same consensus core of cluster 13. (a) Schematics of the secondary structures of cluster 13 sequences identified from SELEX data. Grey bases represent either the invariant universal stem sequence or sequences where we see no base consensus among the SELEX hits in cluster 13. Free bases in the junction core are coloured in red while the paired bases in the core are coloured in blue. Bases coloured in maroon in the ‘Loop1’ region are bases where we see the same consensus among all sequences in the cluster. (b) DMS data of two members (CORT2 and CORT3) of cluster 13. Single mutation data from each are shown, and a black arrow points to the one base in ‘Loop1’ where we see no consensus among SELEX hits. The double mutation data shown are used to infer base-pairing as described in Fig 3c, where blue shows cases where the function of the SSA is rescued preferentially by other Watson-Crick base pairs. A black outline of each cell denotes regions of base-pairing predicted by Mfold. The secondary structure schematics of CORT2 and CORT3 are based on Mfold predictions. Bases predicted to be unpaired are coloured by base essentiality derived from single mutation data (red: low SSA activity, white: high SSA activity). Paired bases are coloured by essentiality of that specific base-pair, as derived from double mutation data (blue: base-pair is not interchangeable and changing to any other base-pairing results in low SSA activity, white: base-pair is interchangeable and similar activity to parental is observed after substitution with any other Watson-Crick base pair. The invariant universal stem is coloured in grey as there is no corresponding DMS data for those bases.

### Deep mutational scanning can identify key drivers of specificity

The cortisol SSAs selected by Yang *et al.*^6^ and the ones we describe above were both selected from the same N30 library. A key difference in how they were selected is that the Yang *et al.* SSA selection protocol used multiple rounds of counterselection to ensure that the cortisol SSAs would only respond to cortisol but not to highly related steroid hormones (Supp Fig 6)— we did not do this since we wanted to capture a more diverse set of cortisol binders without counterselection. The result is that while both CSS.1 and our selected CORT1 are both excellent cortisol sensors (and in fact CORT1 is substantially more sensitive (Fig 10a)), CSS.1 is far more specific and shows strong ability to discriminate between cortisol, aldosterone and progesterone (Fig 10b) whereas the highly related SSA CORT1 shows only weak preference for cortisol over progesterone and nearly identical response towards aldosterone (Fig 10c). Since our less specific SSAs share a similar structure with CSS.1, we reasoned that this might let us pinpoint features that are key for specificity.

**Figure 10.**
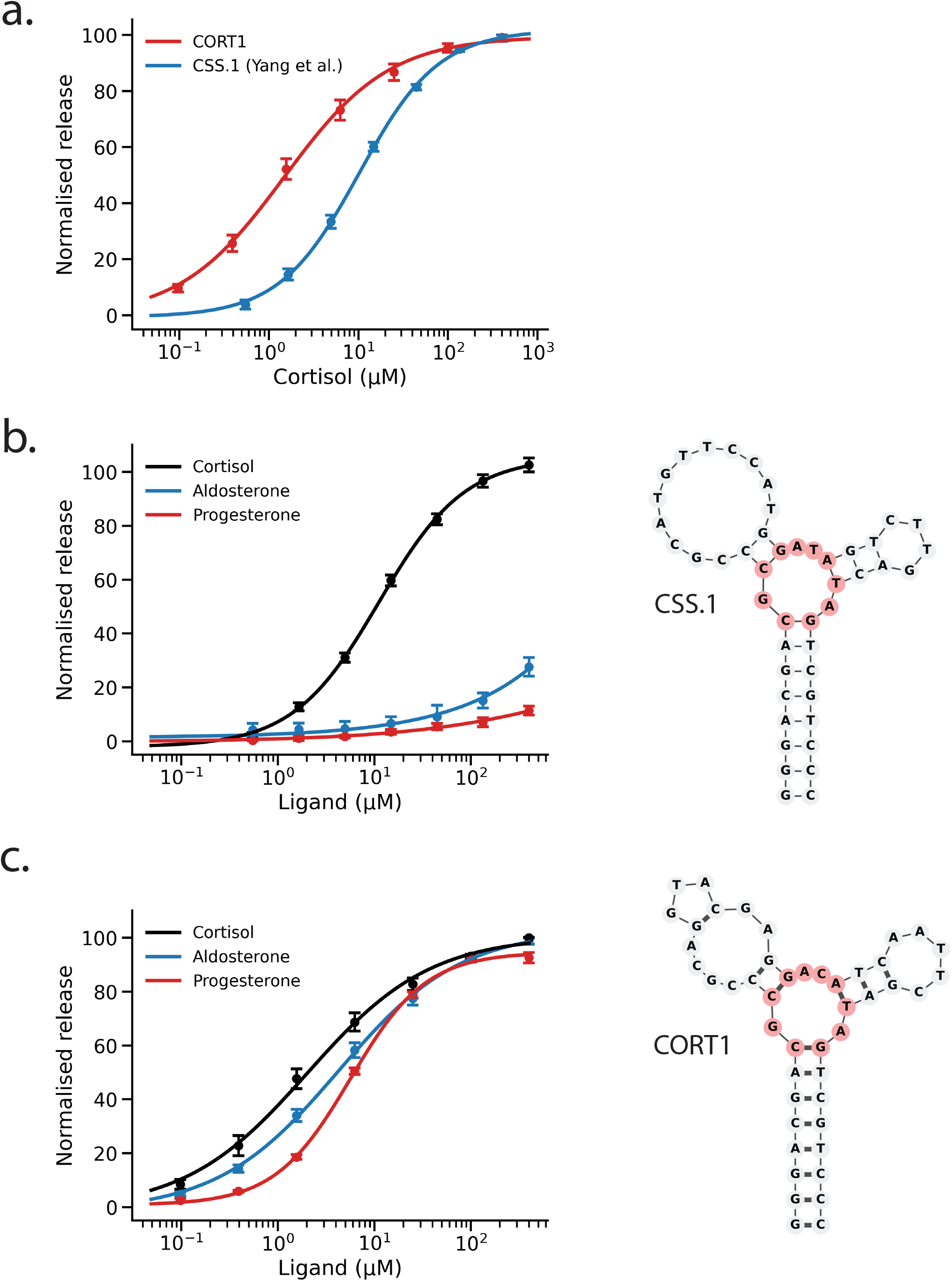
Dose response and specificity of cortisol sensors. (a) Dose response of CORT1 and CSS.1 in the presence of various concentrations of cortisol measured using a fluorescence-based assay where target binding releases a fluorescent oligo (see Material and Methods). (b) Dose response of CSS.1 in the presence of cortisol, aldosterone or progesterone. (c) Dose response of CORT1 in the presence of cortisol, aldosterone or progesterone. In each plot, release is normalised to the minimum and maximum percent release across the concentration range of cortisol for each sensor. Error bars denote standard error from at least 3 independent replicates.

We identified two key essential elements that consistently differ between the specific CSS.1 and the less specific SSAs that we isolated (Fig 11a). The first is a clear difference in the short loop structure that comes off the junction at the core of the SSA — we will call this ‘loop1’ which has different CSS.1 and CORT1 versions (“CSS.1-loop” and “CORT1-loop”; Fig 11a). The sensors we identified by SELEX have a strong consensus across that CORT1-loop and those highly conserved bases are essential for activity based on deep mutational scanning (Fig 7b, Supp Fig 7), yet that entire loop is very different in sequence to the corresponding essential region in CSS.1. Thus while both CORT1 and CSS.1 have essential loop1 regions, these regions are completely different — what does this loop1 region do? To investigate this, we replaced the CSS.1-loop with the CORT1-loop and tested if this hybrid SSA still responds to cortisol and if it can still discriminate between cortisol and progesterone. As shown in Fig 11b, the hybrid CSS.1-CORT1-loop SSA can still respond well to cortisol, albeit with a slightly lower EC50 as CSS.1 itself, indicating that the CSS.1-loop and CORT1-loop are relatively interchangeable in terms of activity despite having little sequence similarity. However, they show a clear difference — the CSS.1-CORT1-loop hybrid can no longer discriminate between cortisol and progesterone, indicating that this ‘loop1’ region is critical for specificity. The CSS.1 version is specific, and the CORT1 version is not — this analysis has thus separated SSA ‘sensitivity’ and ‘specificity’ into two discrete regions of the SSA.

**Figure 11.**
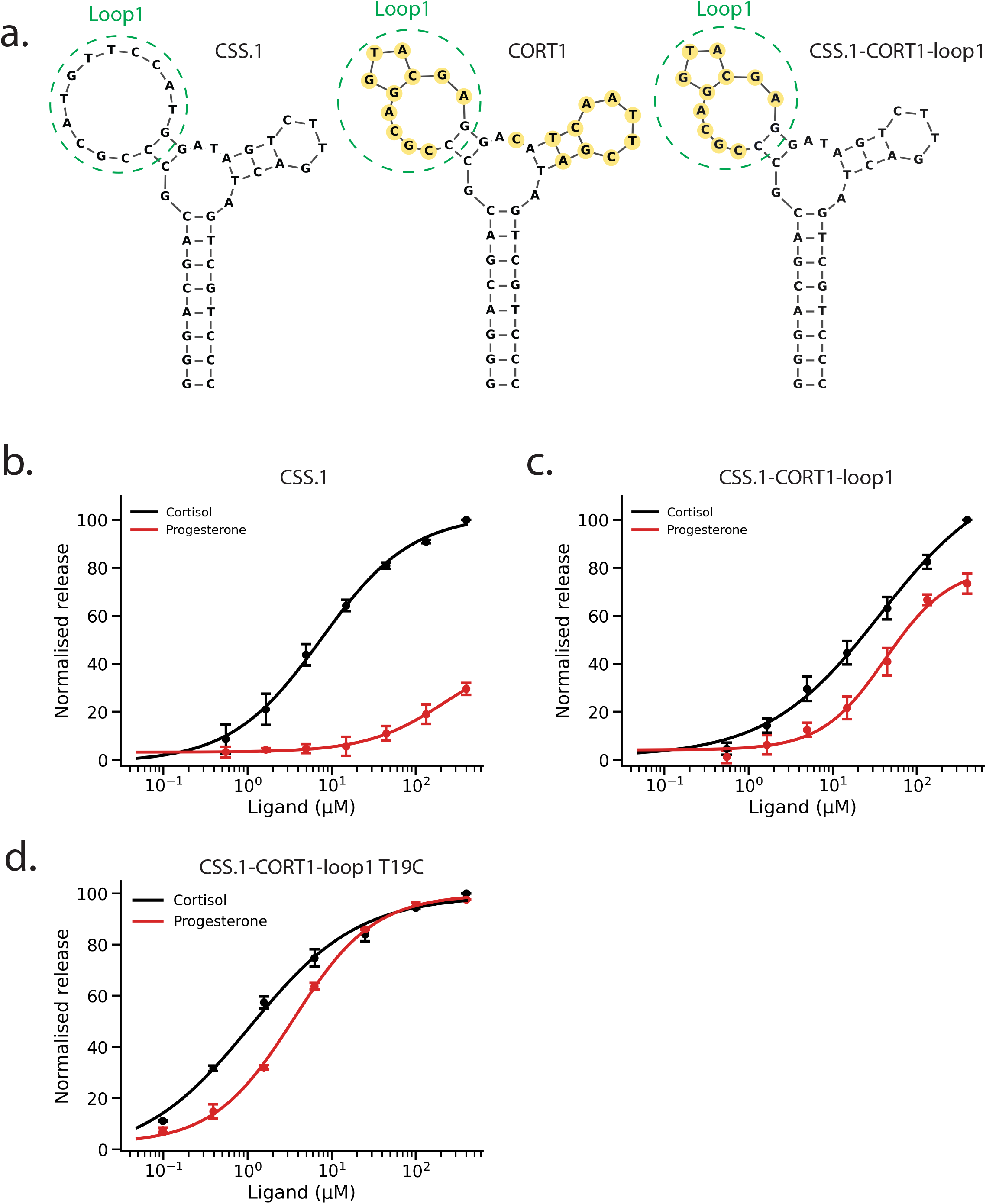
The ‘loop1’ regions of CSS.1 and CORT1 are involved in target specificity. (a) Schematics comparing the secondary structures of CSS.1, CORT1 and the CSS.1-CORT1loop1 hybrid. The regions that differ between CORT1 and CSS.1 and between CSS.1-CORT1-loop and CSS.1 are highlighted in yellow in the respective structures. (b-d) Dose response of SSAs measured using a fluorescence-based assay to measure release of a fluorescent oligo on target binding. (b) High target specificity is observed with CSS.1 in the presence of either cortisol or progesterone. (c) Limited selectivity of the CSS.1-CORT1-loop1 hybrid when tested in the presence of either cortisol or progesterone. (d) The T19C mutant of the CSS.1-CORT1-loop1 hybrid shows increased sensitivity in cortisol (and progesterone) at levels comparable to the CORT1 SSA. In each plot, release is normalised to the minimum and maximum percent release across the concentration range of cortisol for each sensor. Error bars denote standard error from at least 3 independent replicates.

The second key difference is a single base in the core of the 3-way junction (Fig 11a) — CSS.1 has a T whereas all the new sensors in the CORT1 cluster have a C. We find that switching this base in either sensor disrupts sensor activity i.e. a C19T CORT1 is inactive, a T19C CSS.1 is inactive and thus the context of this base clearly matters (Fig 2a, Supp Fig 7a). Intriguingly, we also find that there is some functional interaction between the core of the sensors and the loop1 region — while a T19C CSS.1 is inactive, a T19C CSS.1-CORT1 loop1 hybrid is functional and in fact has increased sensitivity comparable to the CORT1 sensor (Fig 11c). Thus T19 is compatible with a CORT1 loop1 and a C19 is compatible with a CSS.1 loop1, but other combinations (like the T19 CSS.1 hybrid) are non-functional. It is unclear what is the mechanistic basis for this interaction between loop1 and position 19.

Our data above suggest that ‘loop1’ somehow determines specificity — which specific bases in ‘loop1’ do this? To drill down further into which bases in the ‘loop1’ region affect specificity, we used two approaches — both suggested that the same 3 bases are responsible for specificity. Firstly, we noticed that the specific CSS.2 and CSS.3 sensors are strikingly similar in the ‘loop1’ region after we re-aligned those sequences based on our new structure predictions (Fig 5c). Apart from 1 base in the ‘loop1’ region that we find non-essential in our DMS data, all other bases in the ‘loop1’ region share 100% identity between the two sensors, suggesting they might be important for function. We then aligned our CORT1 sensor to these 2 sequences and noticed that among essential bases there were only 3 bases that differed between our non-specific sensor and the CSS.2 and CSS.3 sensor in this conserved region (Fig 12a). Since the ‘loop1’ region is critical for specificity, and since CSS.2 and CSS.3 are specific and share these bases and CORT1 is non-specific and has different sequences there, this suggested that these might be key for specificity.

**Figure 12.**
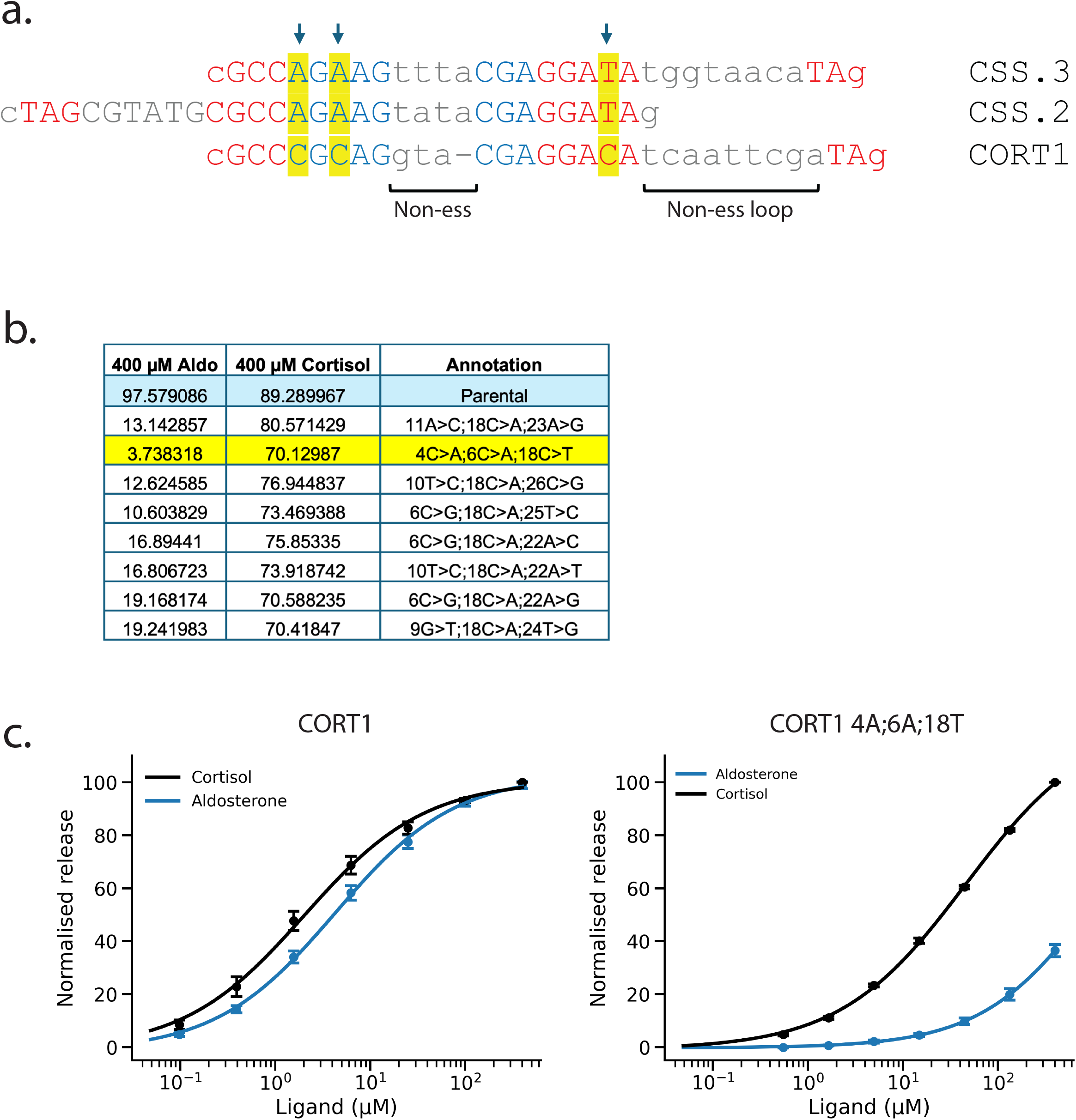
CORT1 specificity is increased by three base changes. (a) The sequence of CORT1 is compared to the sequences of CSS.2 and CSS.3 by aligning them by their conserved core elements (red text). Blue text indicates essential bases (derived from DMS data) in the ‘Loop1’ region of these SSAs. Non-essential elements are coloured in grey and in lowercase. Yellow highlighted bases show the three bases that differ between CSS.2-3 and CORT1 among the essential bases. (b) The top 8 hits from DMS data that show increased difference in SSA release in the presence of cortisol versus aldosterone. Hits are ordered by the difference in release observed between cortisol and aldosterone. The release values in the parental CORT1 sequence are highlighted in blue in the table, and yellow highlights the triple mutant with increased selectivity, that are the same 3 bases as identified in (a). (c) Dose response of CORT1 and the triple mutant are measured in the presence of cortisol or aldosterone using the same fluorescence-based assay as described above. In each plot, release is normalised to the minimum and maximum percent release across the concentration range of cortisol for each sensor. Error bars denote standard error from at least 3 independent replicates.

Secondly, we performed DMS of our CORT1 sensor with aldosterone instead of cortisol as a target — our CORT1 sensor cannot distinguish between cortisol and aldosterone (Fig 10c) and we reasoned that DMS could identify mutations that increase sensor specificity. We identified several mutant CORT1 sequences that now show strong specificity for cortisol over aldosterone — the second highest hit had mutations in the same 3 bases we identified by sequence comparisons between non-specific CORT1 and specific CSS.2 and CSS.3 (Fig 12b). Since both sequence comparisons and DMS converged on these 3 bases as a possible driver of specificity, we synthesised a version of CORT1 that contains these 3 changes — the mutant CORT1 shows marked selectivity between cortisol and aldosterone (Fig 12c). We conclude that DMS can not only be used to uncover the structural basis for SSA activity, it can also pinpoint the key sequences that drive target specificity.

### Deep mutational scanning can select variants with altered target specificity

Cortisol and progesterone are similar target molecules but CSS.1 only responds to cortisol but not progesterone — can we use DMS to identify mutant versions of CSS.1 that have reversed specificity i.e. that now respond to progesterone and not cortisol? We did deep mutational scanning on CSS.1 and compared the ability of every mutant in the pool to respond to cortisol and to progesterone. We identify 2 double mutants that could now preferentially detect progesterone over cortisol (Fig 13a-b). Both mutants are predicted to make an additional base-pair in the core of the 3 way junction and a further base-pair extending out of the junction, thus reducing the size of the central pocket (Fig 13c). This might explain the greatly reduced ability to detect cortisol which has multiple additional side groups compared with progesterone.

**Figure 13.**
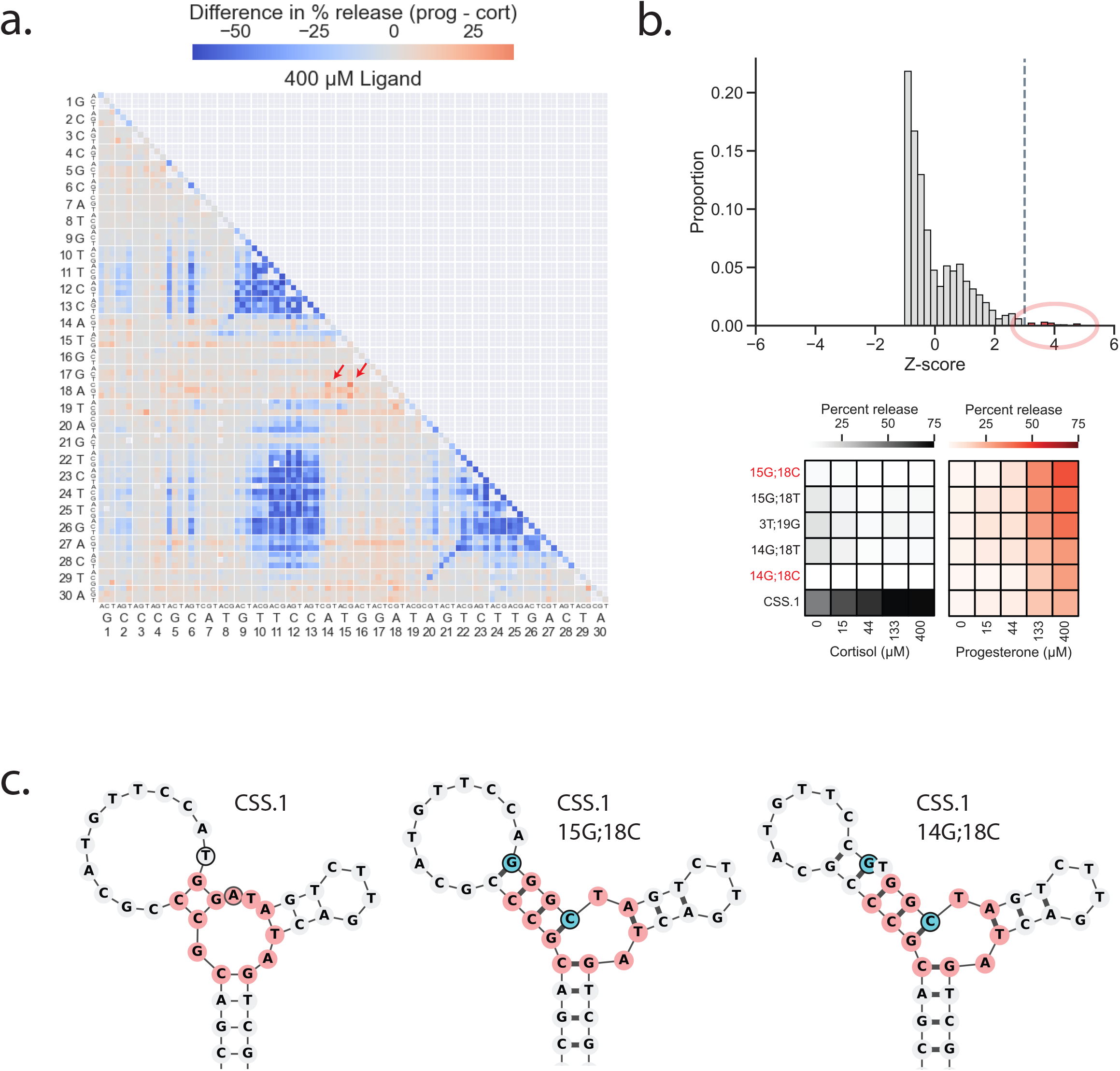
DMS can identify variants with altered target specificity. (a) The DMS double mutation profiles of CSS.1 when selected against both cortisol and progesterone. The difference in release when incubated in 400 µM progesterone relative to 400 µM cortisol is shown. Red show greater release in progesterone in the double mutant relative to cortisol and blue show greater release in cortisol. Arrows point to the 15G;18C and 14G;18C double mutants. (b) The relative performance of all functional double mutants in progesterone relative to cortisol are used to calculate z-scores. All z-scores > 3 (increased activity of the double mutant in progesterone relative to cortisol) are highlighted in the plot. Heatmaps are shown from the top 5 hits in this set across a range of doses tested in cortisol and progesterone. The parental CSS.1 sequence is included for comparison. (c) Predicted secondary structures of two of the top hits from the DMS screen compared to CSS.1. Red is used to highlight the conserved 3-way junction core, and blue highlights the double changes in each structure, both predicted to result in 2 additional base pairs.

To better understand the structural basis for the double mutant CSS.1 15T>G;18A>C SSA that shows a change in specificity, we carried out DMS to identify the base-pairing and base requirement in this CSS.1-derived SSA and found that there has been a large rewiring of base-pairing in this double mutant SSA that confirms Mfold predicted structure: around half of the base-pairs have changed relative to the parental CSS.1and many positions that were previously essential now appeared to have little constraint, including in the ‘loop1’ region that we found essential for cortisol specificity (Fig 14a-b).

**Figure 14.**
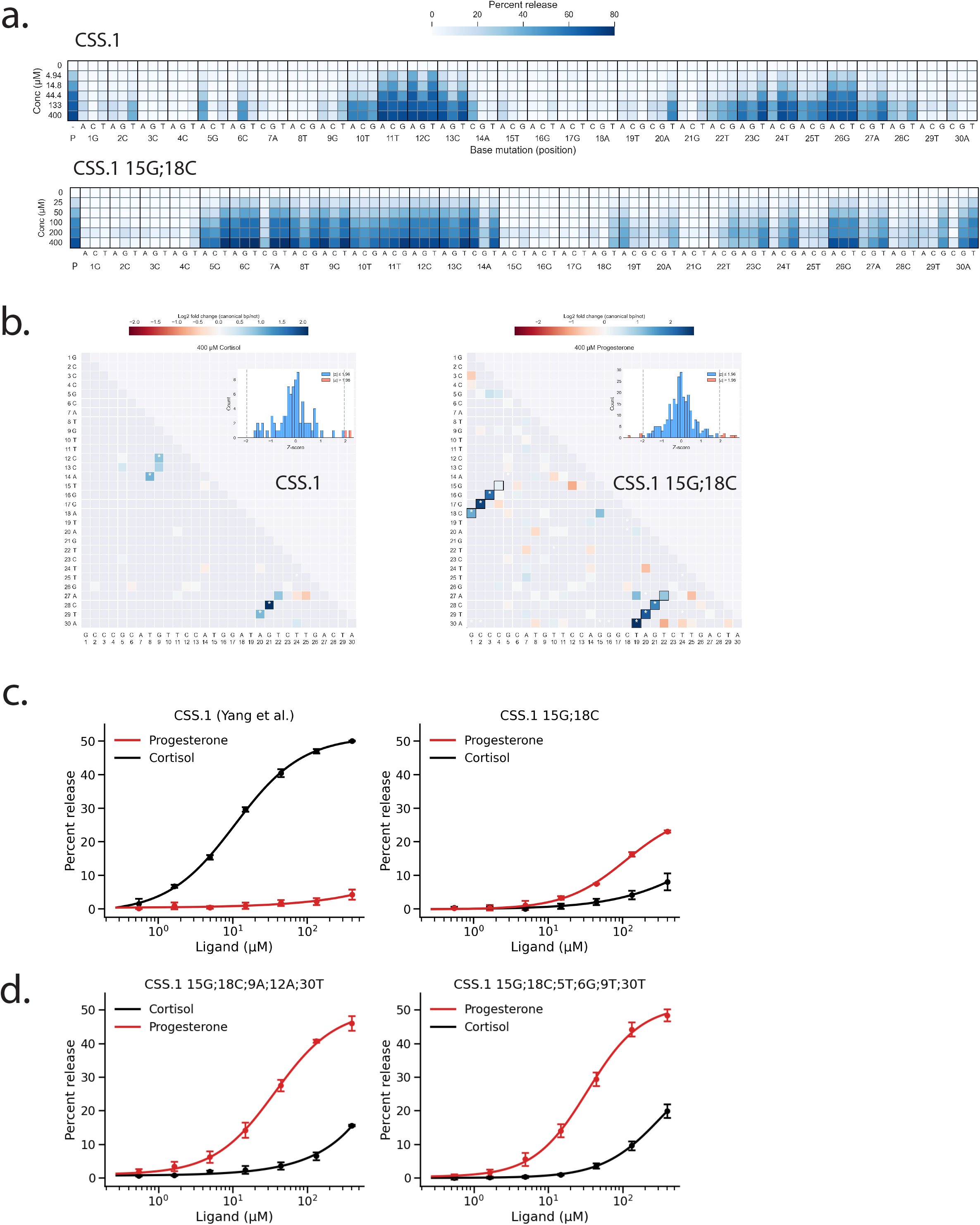
DMS data suggest structural re-wiring in the CSS.1 15G;18C double mutant. (a) Single mutation plots of CSS.1 and the 15G;18C double mutant. DMS was carried out on the CSS.1 sequence in various concentrations of cortisol while DMS was carried out on the double mutant in various concentrations of progesterone (with a narrower concentration range). (b) Base-pairing patterns inferred from double mutation profiles in CSS.1 and the 15G;18C double mutant at 400 µM cortisol and progesterone respectively. Blue shows cases where the function of the SSA is rescued preferentially by other Watson-Crick base pairs at those two positions (method described in Fig 3c). (c) Dose response of CSS.1 and the double mutant are measured in the presence of cortisol or progesterone using the same fluorescence-based assay as described above. (d) Dose response of the 5x and 6x mutants of CSS.1 that show increased response in the presence of progesterone. Error bars denote standard error from 3 independent replicates.

However, while the double mutant CSS1 15T>G;18A>C SSA shows a change in specificity compared with the parental CSS.1, it still only responds poorly to progesterone with a EC50 >100 μM (Fig 14c). We thus did an additional round of DMS using the 15G;18C double mutant as the starting sequence to try to isolate additional mutations that could improve the response of this double mutant to progesterone. This yielded multiple SSAs that differ from the starting CSS.1 by a total of 5-6 independent mutations and that now detect progesterone far better than cortisol with much increased sensitivity (Fig 14d).

We conclude that DMS can not only identify many of the base-pairing interactions and base requirements in a specific SSA, but can also be used to change target specificity and gain insights into how an SSA achieves specificity.

## Discussion

AlphaFold has transformed protein structure prediction — it is possible to predict the structure of almost any protein to near atomic resolution within seconds from its primary amino acid sequence^13^. However, for structured nucleic acids like aptamers, ribozymes, and riboswitches, structure prediction is still very challenging^18,19^. In part this is because many of these nucleic acid machineries undergo major conformational changes on target binding, but in large part this is because there is too little structural data. AlphaFold trained on over 200k solved protein structures and deep multiple sequence alignments — there are no analogous rich datasets for nucleic acids^19^. How can we will this knowledge gap? It is unlikely that there will be tens of thousands of solved structures for structured nucleic acids bound to their targets in the foreseeable future — in this study we wanted to try to use Deep Mutational Scanning (DMS) as a rapid method to capture insights into aptamer structure.

DMS has been used to great success in protein structure analysis^14,15^. The most basic data are the effects of every possible single amino substitution on protein stability or activity and these data can be very valuable for interpreting how inherited mutations may impact protein activity. However, for structural insights, the key is double mutations — these can reveal functional interactions between amino acids and these can be used as constraints in structure predictions. We took a similar approach here.

We used DMS to investigate the structure-function relationships in 3 published structure-switching aptamers (SSAs) that all specifically recognise cortisol^6^. Two of these were thought to have a highly related functional core (CSS.1 and CSS.3), whereas the third, CSS.2, is predicted to have a very different structure. We measure the effect of every single mutation, every double mutation, and every single base on activity. Just as with protein DMS, the double mutation data reveals base-base interactions and this gives key insights into how these SSAs are structured and how they recognise their targets.

First, our DMS data show the published and predicted base-pairing in CSS.1 and CSS.3 are likely correct but the predicted structure of CSS.2 is likely wrong — we cannot confirm any of the predicted base-pairs and instead identify a different set. Second, we find that using our DMS-derived base-pairings indicates that CSS2 likely forms a structure that has an identical core to CSS.1 and CSS.3 — our DMS data thus suggest that all three published cortisol sensors likely bind their target in a very similar way.

CSS.1, CSS.2, and CSS.3 all appear to recognise cortisol via a similar structural mechanism. All three were selected using SELEX and although many thousands of sequences were likely enriched during SELEX, very few were sequenced, and we thus wanted to explore more deeply how DNA-based SSAs might recognise cortisol. We thus carried out SELEX from an identical library to that used for selection of CSS.1-CSS.3 and identified many more sequences that could recognise cortisol. The highest ranked structural family we identified resembled the CSS1-like core but every one of our new set of SSAs also had an additional highly conserved loop (‘CORT1 loop1’) that was not found in CSS.1-3. CSS.1-3 were not only selected to bind cortisol — they were also counter-selected to ensure that they did not bind related molecules and we reasoned that this difference between our new SSAs and CSS.1-3 might affect specificity. We confirm that CSS.1 only responds to cortisol but not progesterone or aldosterone but that CORT (a representative of our new SSAs) was an excellent cortisol sensor but showed little specificity for cortisol over progesterone or aldosterone. We could tie this to the CORT1 ‘loop1’ region — while a hybrid CSS1 that included the CORT1 loop1 region was still an excellent cortisol sensor, it was no longer specific. By combining assays on hybrid molecules between CORT1 and CSS.1 and DMS to study positions that affected specificity, we identify three critical bases that were conserved in CSS.2 and CSS.3 and showed that engineering these three base changes in CORT1 now made it into a specific cortisol SSA. These data show that at least for these related cortisol SSAs, the regions responsible for response to the target and the regions responsible for target specificity are distinct and that there are complex functional interactions and compatibilities between these regions.

DMS had helped us uncover the determinants of specificity in the CSS.1-3 family of specific cortisol sensors — we also used it to identify mutants of CSS1 that had altered specificity. The CSS.1 parent recognises cortisol but does not respond to progesterone — we found that a double mutant of CSS.1 now preferred progesterone to cortisol and identified multiple quintuple mutants of CSS.1 derived from that double mutant that are excellent progesterone and not cortisol sensors. What was surprising to us is the double (and 5X) mutant of CSS.1 showed a dramatic rewiring of base-pairing: over half the base-pairings were different in the double mutant than in the CSS.1 parent.

Taken together we thus conclude that DMS is a powerful and rapid way to identify base-pairing interactions that underlie the structures of SSAs when they bind their targets. While this may not capture every base-base interaction, the DMS base-pairings can be very powerful in testing structural models and the type of structure-function data generated by DMS is very rich and powerful for modeling. We note that base-pairing constraints derived from chemical accessibility have yielded in silico models of RNA structures that outperform AlphaFold^20^. We believe DMS will also be a rapid and rich source of constraints for in silico modeling of SSA structures and generating DMS data for many hundreds of aptamers may similarly guide in silico models of nucleic acids structures.

## Materials and Methods

### Reagents

All oligonucleotides were synthesized by Integrated DNA Technologies (IDT) and dissolved in nuclease-free water at a concentration of 100 µM. The CSS.1-3 sequences used were adapted from Yang *et al.*^6^ All oligonucleotide sequences used for SSAs are listed in Supplementary Table 1. Stock solutions of cortisol, progesterone and aldosterone (Sigma) were made in 100% ethanol with a final working concentration of 5% ethanol. All binding assays were done in PBS (pH 7.4, Gibco) + 5 mM MgCl_2_ unless otherwise noted.

### Ligand binding assay

For DMS, SELEX and individual dose response assays, we use the same ligand binding and oligo release method as outlined below with the following difference: for DMS and SELEX assays we bind the LBOs to an immobilized short oligo, whereas for individual dose response assays, we bind an SRO to immobilized LBOs (as described previously^11^).

For each sample concentration, 2.5 µL of Dynabeads MyOne Streptavidin C1 magnetic beads (Invitrogen) were washed 3 times in Bind & Wash buffer (5 mM Tris-HCl (pH 7.5), 0.5 mM EDTA, and 1.0 M NaCl) as per manufacturer’s protocol and finally resuspended in SELEX buffer (1M NaCl, 20 mM HEPES (pH 7.5), 10 mM MgCl_2_, 5 mM KCl) at 2x initial bead volume. 25 pmol of LBO and 125 pmol of SRO (5 μL total for each sample concentration, diluted in SELEX buffer) were heated at 95°C for 5 min and slowly cooled to 25°C. The oligos were then added to the resuspended beads at equal volume and incubated on a rotator for 30 min at room temperature.

The beads were then washed 2 times in PBS + 5 mM MgCl_2_ buffer and resuspended in 25 μL of the same buffer and incubated for another 40 min. The beads were then washed once and resuspended with various concentrations of ligand. The samples were incubated for 45 min on a rotator at room temperature and the supernatant was collected (‘supernatant’ sample). For DMS and dose response assays, the beads were also then resuspended in 25 μL of PBS without magnesium and heated at 95°C for 4 min to release the remaining bound SROs, and the supernatant sample collected (‘remainder’ sample).

### Fluorescence readout

For single dose response assays for individual sensors, we use a fluorescence readout as described previously^11^. In outline, each SSA consists of two oligos – a biotinylated Ligand-Binding Oligo (LBO), which is attached to streptavidin magnetic beads, and a Short Release Oligo (SRO) with a 6-FAM fluorophore which base pairs with the LBO. A conformational change on target binding induces the release of the corresponding SRO, and levels of the released SROs are read out using a plate reader.

The fluorescence intensity for each sample was then measured using a FLUOstar Omega microplate reader (excitation 485 nm, emission 520 nm) and percent release is calculated as: supernatant RFU/(supernatant RFU+remainder RFU)_x 100. For normalized release, the minimum release is normalized to 0 and the maximal release to 100. For dose response curves, a four-parameter logistic (4PL) model was then used for curve fitting.

### Deep mutational scanning

‘Doped synthesis’ oligos were ordered from IDT using a 94:2:2:2 mixture of bases at each position of the 30-mer variable region. Oligos were ordered with a common stem region and PCR flanks as described for SELEX experiments in Yang *et al.*^6^ LBOs were annealed to immobilized short oligos at a 1:3 ratio. Ligand binding to each pool of oligos was assayed at 6-8 target concentrations as described above and 2 additional ‘boil off’ samples (‘input’) were collected to estimate input amounts of LBOs that were bound to the immobilized short oligos before any ligand-mediated release.

PCR was then carried out on the input and release samples using unique barcoded forward and reverse primers for each sample condition at the same but limited number of cycles to limit saturation (∼10-14 cycles, determined empirically for each SSA). Samples were amplified using 2x KAPA HiFi HotStart ReadyMix (Kapa Biosystems) in 25 μL reactions (+2 μL DNA, 3 μL of F and R primers at 10 μM each). Primers used at this step contains unique sample barcodes, diversity spacers and partial P5/P7 Illumina adapter sequences for sequencing library preparation.

For each SSA-ligand combination, the PCR reactions were then pooled and cleaned up using Monarch Spin PCR & DNA Cleanup Kit (New England Biolabs). The eluate was then diluted to ∼5 ng/μL and used in a second round of PCR to add the i5/i7 index sequences and full P5/P7 adapter sequences. A 30 μL KAPA HiFi PCR reaction was carried out at this step with 2 μL DNA and 1.5 μL 10 μM primers and run at 8 cycles. Libraries were then cleaned up using Monarch Spin PCR & DNA Cleanup Kit, pooled, and a size selection step was performed using NucleoMag NGS Clean-up and Size Select beads (Macherey-Nagel) at a 1.2x bead to sample. The samples were then resuspended in 10 mM Tris-HCl, pH 8.0 and sent for Illumina sequencing on a NovaSeq X at The Centre for Applied Genomics (Toronto, CA).

The raw sequencing data was demultiplexed and the variable 30mer region parsed using AptaSuite^21^. The collated counts for each unique 30mer were then used with custom Python code to estimate relative percent release values for each sequence. Since the input and release fractions were amplified and sequenced at different depths – to normalize values we applied a universal size factor to counts in each input and release sample so that the estimated percent release values fall between 0 and 100, at levels that match the actual percent release we observe with the parental sequence in our fluorescence assays. We note that the percent release calculated in this manner are not absolute percent release values, but rather relative values that allow us to compare between sequences in each DMS experiment (since each sequence in the pool is multiplied by the same size factor) but not directly comparable between experiments. All data were then analysed and plotted using Python.

### SELEX

For SELEX experiments, we use the same N30 SSA library design, capture oligo, and primers as described by Yang *et al.*^6^ and follow a similar protocol with some modifications. For the first round of SELEX, for each target selection 200 pmol of the starting N30 library was added to 5x excess capture oligo in SELEX buffer. The samples were then heated at 95°C for 5 min and slowly cooled to 25°C. The annealed and folded oligos were then added to 100 μL of Dynabeads MyOne Streptavidin C1 magnetic beads and incubated on a rotator at room temperature for 30 min (total 400 μL volume). Beads were then washed in PBS + 5 mM MgCl_2_ buffer as described above and further incubated with the same buffer for 10 min. Samples were then split into 4 tubes (100 μL each), resuspended with buffer only (with 5% ethanol) for 5 min, after which sample was replaced by 100 μM cortisol and incubated for 10 min. The supernatant was then collected and pooled and then washed and concentrated to a final volume of ∼50-100 μL using a 3-kDa Amicon Ultra-0.5 centrifugal filter (Millipore).

A small-scale PCR (5 μL DNA in a 50 μL Taq PCR reaction) was then performed with variable number of cycles and run out on an agarose gel to determine the number of cycles to use for later large-scale PCR to avoid saturation. A larger scale PCR reaction (1 mL initially and 600 μL in later rounds) was then carried out to amplify the library for the next round. The amplified products were then cleaned up and concentrated using Monarch Spin PCR & DNA Cleanup Kit and eluted in H_2_O.

Preferential strand digestion by lambda exonuclease was used for the strand separation step. Specifically, for the PCR reaction a forward primer with 6 phosphorothioate bonds at the 5’ end was used to protect the desired forward strand from digestion by lambda exonuclease, and a 5’ phosphorylated reverse primer was used to promote digestion of the other strand. After PCR cleanup, 5 μL of the purified library was kept for sequencing later, and the rest were incubated with lambda exonuclease (NEB) for 30 min, followed by heat inactivation at 75° for 15 min. The salt composition and pH of the sample was then re-adjusted by adding 1/5 volume of concentrated SELEX buffer. 5x excess capture oligo was then added to this mix and heated at 95°C for 5 min and cooled to 25°C to repeat the cycle. ∼50-100 pmol of the library was carried forward to the next round of SELEX.

This protocol was then repeated for subsequent rounds with the same steps except with less beads (25 μL) and total volume (100 μL). Rounds 1-5 were selected at 100 μM cortisol (in 5% EtOH), rounds 6-8 at 50 μM cortisol (in 2.5% EtOH) and rounds 9-10 at 25 μM cortisol (in 1.25% EtOH). The TUDCA selection was carried out at these same target concentrations.

Library preparation for sequencing each round was similar to the method described above for the DMS samples. Collected samples from each round were subjected to 2 rounds of PCR to add sample barcodes, diversity spacers and Illumina adapter and index sequences, followed by PCR cleanup and size selection. Samples were then sequenced at ∼20M reads per round on a NovaSeq X at The Centre for Applied Genomics.

### SELEX analysis

The raw sequencing data was demultiplexed and the variable 30mer region parsed using AptaSuite^21^. The collated counts for each unique sequence were then used to identify the top 5000 hits in round 10 of each target selection to re-test. These top 5000 sequences were then reordered in a single pool synthesized by IDT. This library was then assayed and analysed for target binding at various concentrations in the same way as described above for the DMS libraries.

The top re-tested hits were then analysed in two ways. First, we did a simple sequence clustering using FASTAptamer^22^ to identify clusters of sequences using a Levenshtein edit distance of 7. These clusters were then visualized using Cytoscape^23^. Second, for structure clustering, we first used Mfold^24^ to predict the secondary structure of each sequence. We then use the Forgi package^25^ in Python to classify the structural elements from the dot-bracket structures obtained from Mfold. We focus specifically on 3-way junctions in this paper and extract the junction core sequences of each structure that has a 3-way junction using custom Python code. We differentiate and annotate both free and paired bases in each 3-way junction from the 5’ to 3’ end to extract a core motif. To additionally match ‘rotational’ variants, we then ordered each motif alphabetically so that structures with the same core motif in any orientation are clustered together. Due to alternative structure predictions, we find that the same sequence can be sorted into different but related clusters. To prevent over-counting of clusters, we then collapsed these similar clusters together and manually re-assigned some sequences for which we had DMS data to predict the proper cluster assignment (e.g. for CORT1, CORT107).

## Acknowledgements

This research was funded by project grants PJT183712 and PJT204094 to A.G.F. from the Canadian Institute of Health Research (CIHR). We thank Cameron Mackereth and Eric Largy for valuable discussions, members of the Fraser Lab for advice and helpful discussions throughout, Arttu Jolma for guidance on designing sequencing libraries, and The Centre for Applied Genomics (TCAG) for help and advice with sequencing.

## Competing Interests

The authors declare the following competing interests: A.G.F. and J.H.T. are listed as inventors on a provisional patent application filed by the University of Toronto relating to the use of barcoded structure-switching aptamers.

---

**Supplementary Figure 1.**
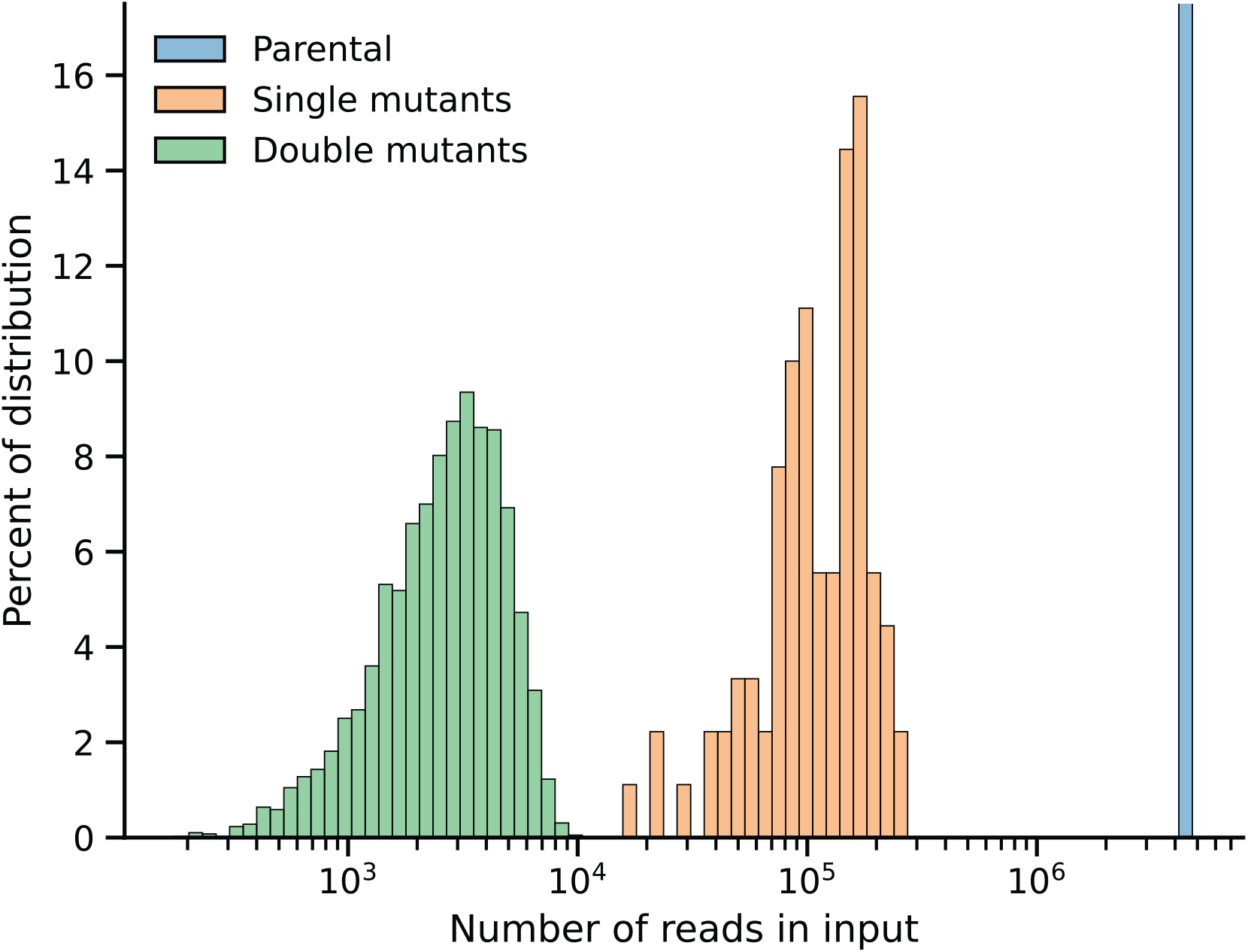
Distribution of read counts of the input amounts of each sequence in the DMS pool before ligand addition. The distribution of the number of reads sequenced for all single and double mutants in the CSS.1 DMS pool are plotted. Double mutants are rarer in the pool than single mutants as expected, though there is large variation between read counts for mutants in each category.

**Supplementary Figure 2.**
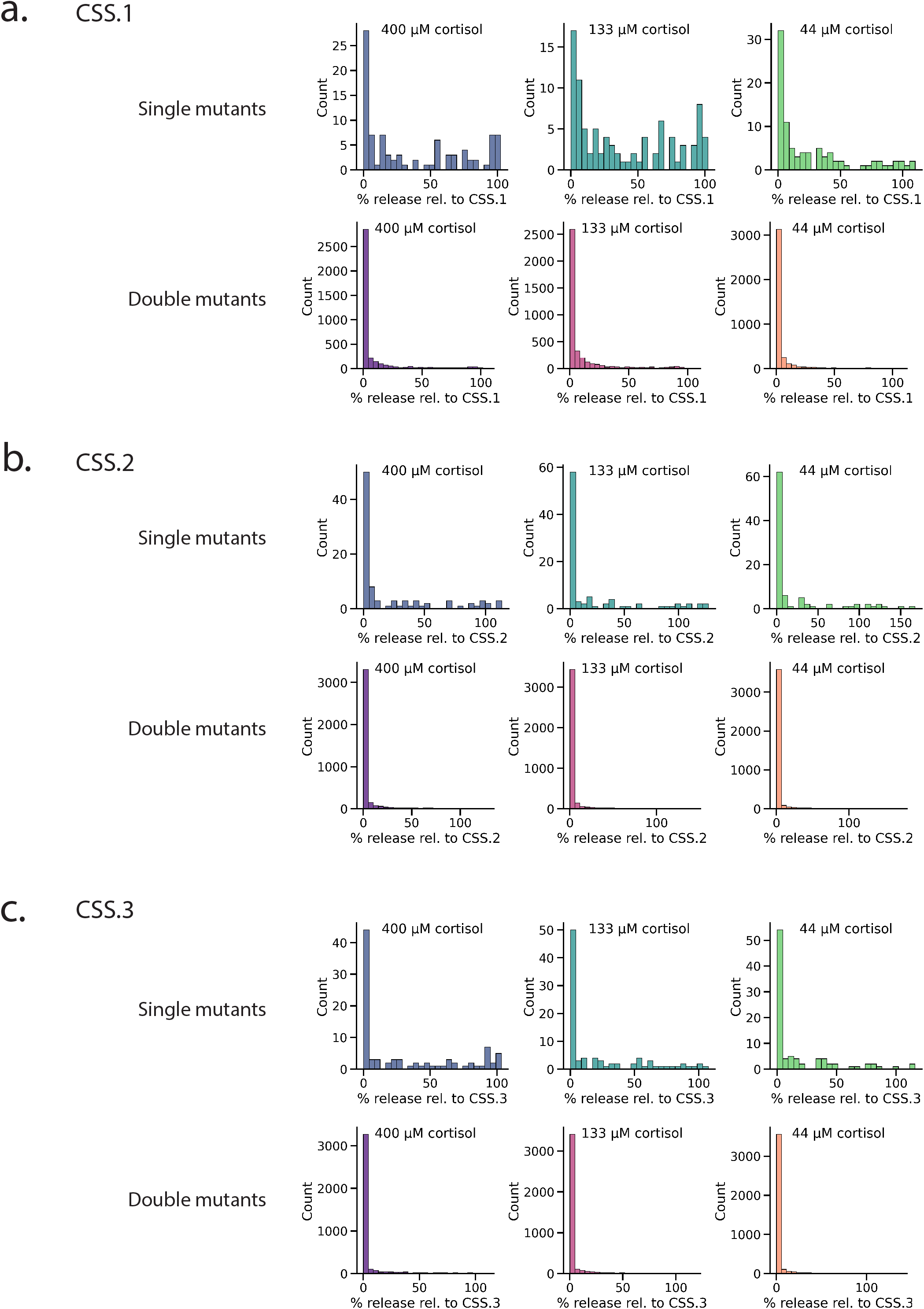
Distribution of release values among all single and double mutants from DMS of cortisol sensors. DMS was carried out in CSS.1 (a), CSS.2 (b) and CSS.3 (c) and the distribution of percent release values in single and double mutants in these screens are plotted at the various concentrations of cortisol tested.

**Supplementary Figure 3.**
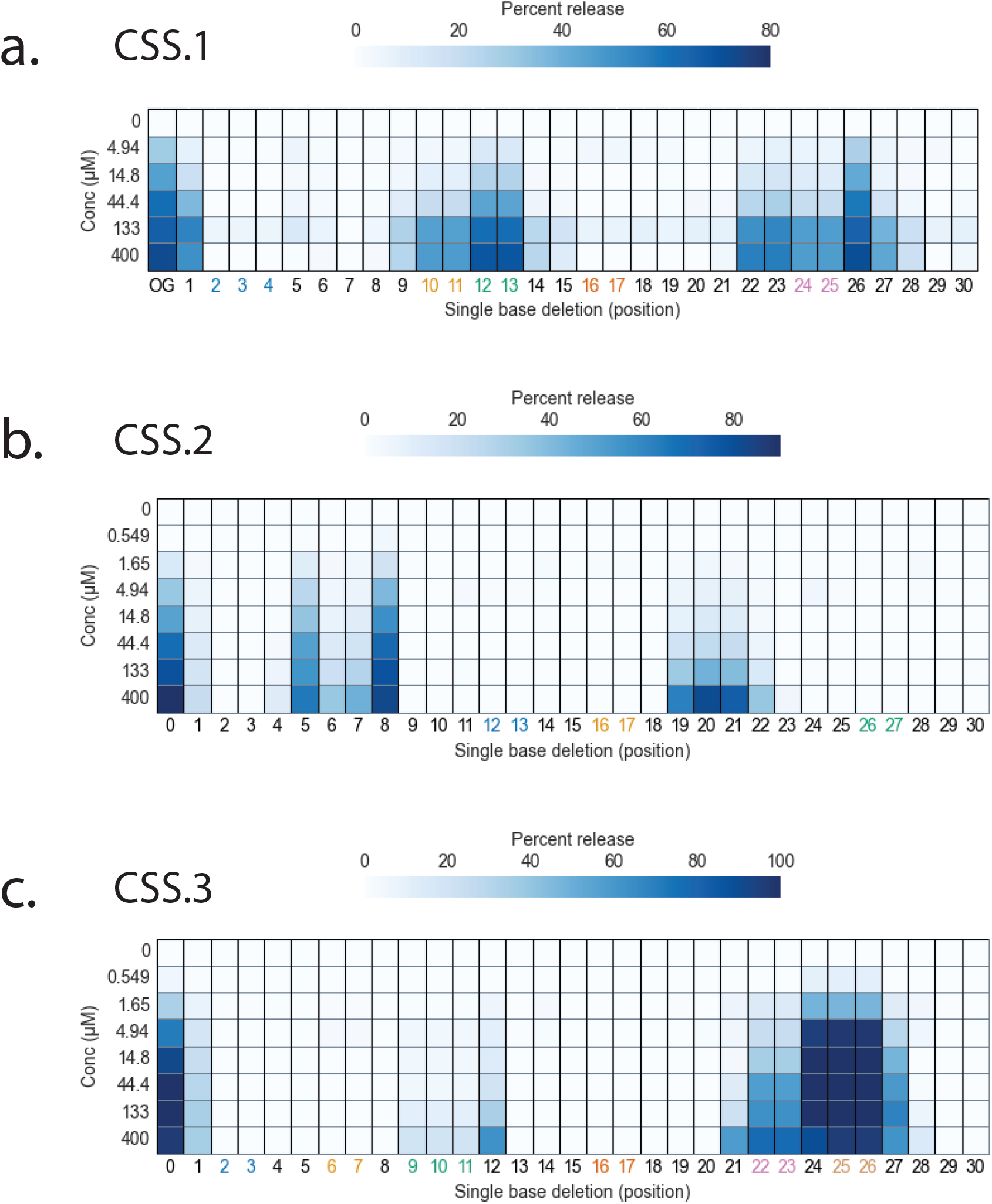
Dose response profiles of single deletion mutants from DMS of CSS.1-3. Heatmaps show the dose response of all single mutants of CSS.1 (a), CSS.2 (b) and CSS.3 (c) across all 30 bases in the variable region. The x-axis shows base position numbers, and numbers that are coloured show positions with the same base at two or more consecutive positions where actual deletion position cannot be differentiated and thus the same response is plotted at each of these consecutive positions.

**Supplementary Figure 4.**
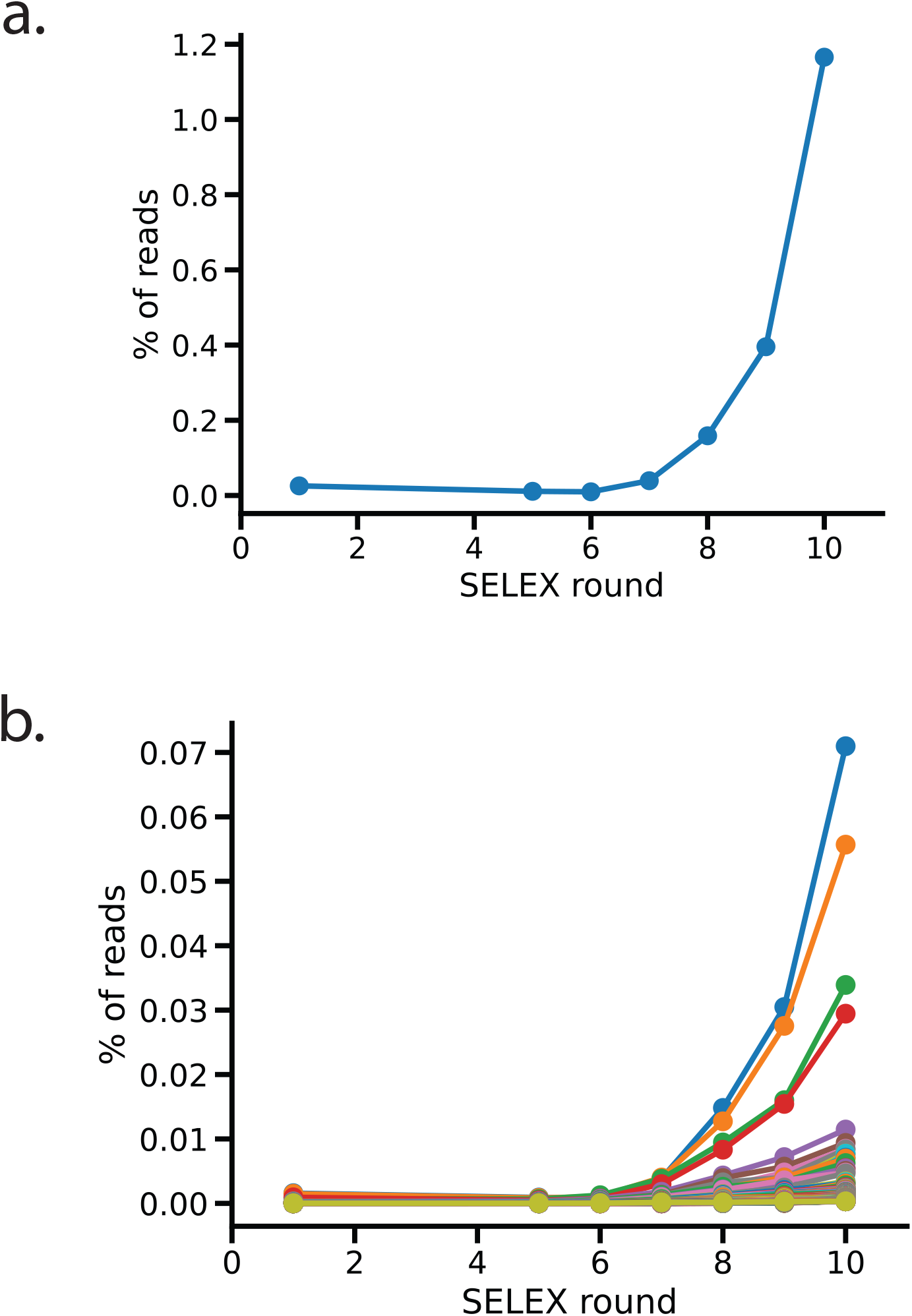
Increase in enrichment of the top 100 SELEX hits across 10 rounds. The SELEX pool was sequenced after round 1 and rounds 5-10 and enrichment of the top 1 sequence (a) and the top 2-100 sequences (b) are plotted. Each sequence count is normalised as a percentage of the total amount of reads sequenced for that round.

**Supplementary Figure 5.**
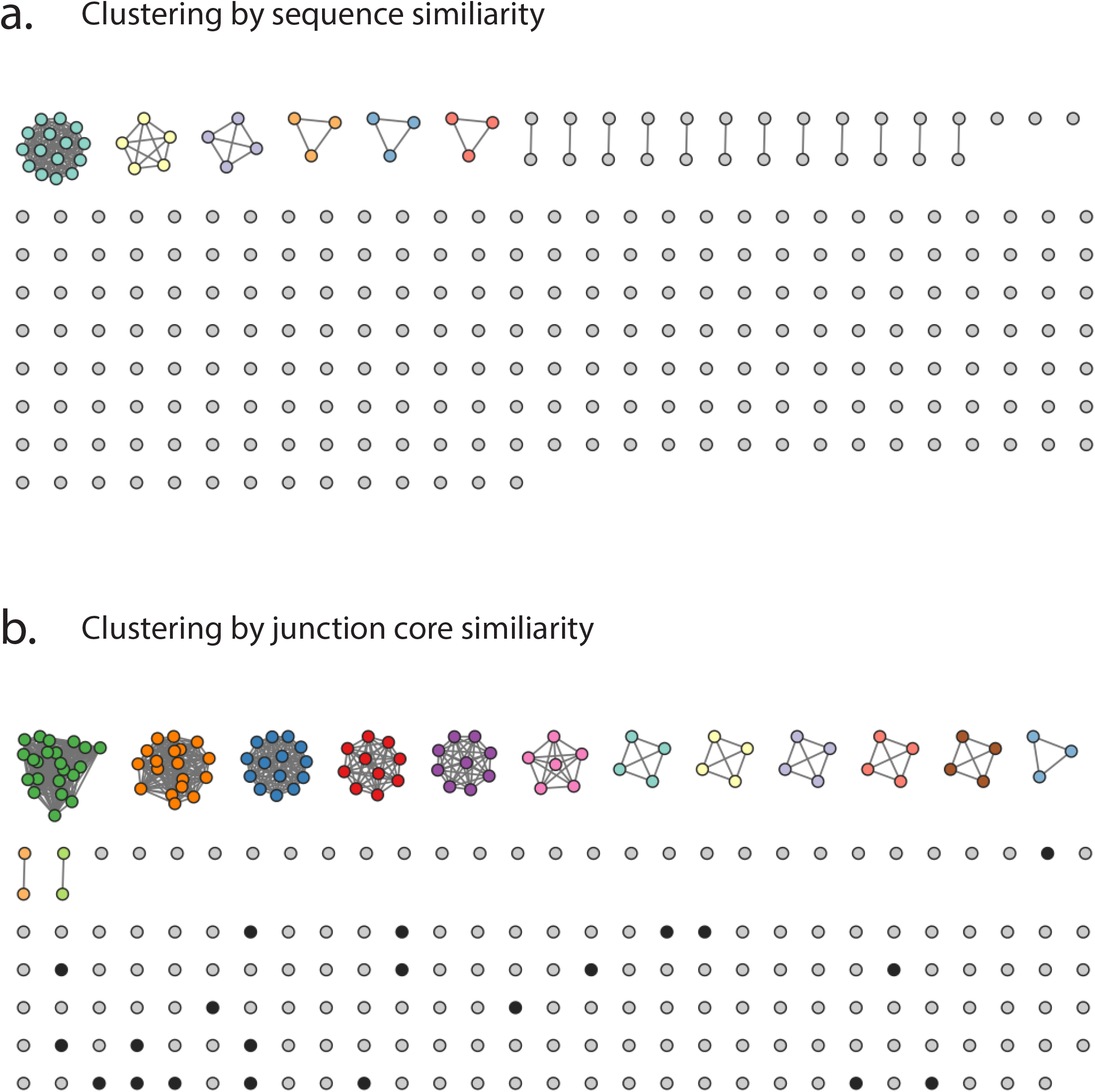
Clustering of the top 277 performing SELEX hits. The top 277 SELEX hits (after pooled re-testing) were clustered either by sequence similarity or by structural core similarity. (a) The sequences were clustered using FASTAptamer, using an edit distance of 7 to define clusters. Cytoscape was used to generate the resulting cluster map. The most populous cluster consists mostly of sequences with a single base change relative to the CORT1 sequence (the most enriched sequence from the SELEX). (b) The sequences were clustered using a custom Python script that identifies the 3-way junction core (see Materials and Methods). All sequences with the same 3-way junction core were clustered together. All coloured or black nodes indicate sequences predicted to form a 3-way junction. Cytoscape was used to generate the resulting cluster map.

**Supplementary Figure 6.**
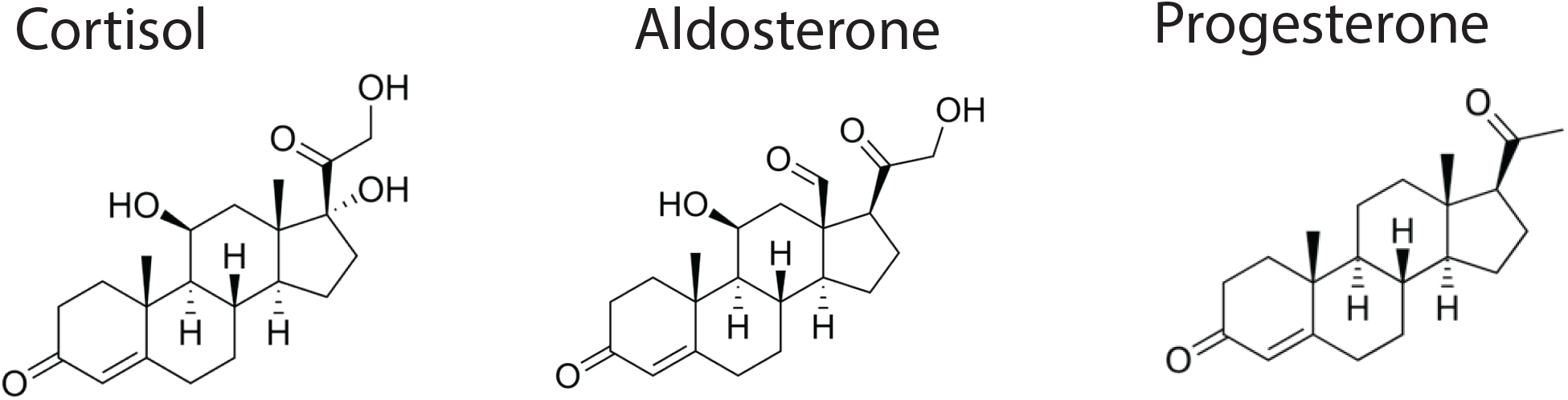
Chemical structures of cortisol, aldosterone and progesterone.

**Supplementary Figure 7.**
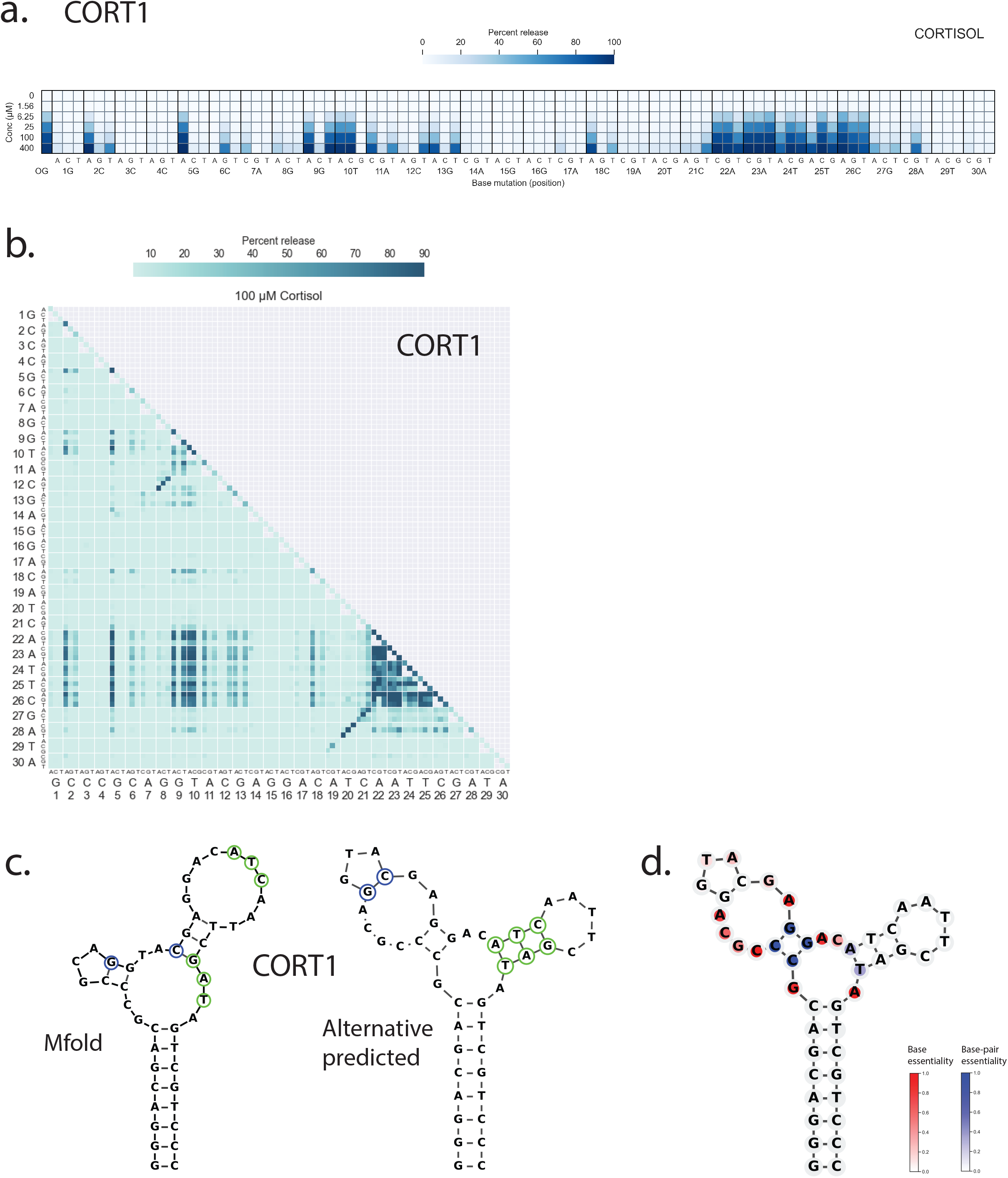
DMS data of CORT1 was used to infer secondary structure different from Mfold prediction. (a) Dose response of all single mutants of CORT1 in the presence of various concentrations of cortisol. A darker blue indicates higher levels of release. (b) Percent release of all double mutants of CORT1 in the presence of 100 µM cortisol. Darker shades indicate higher levels of SSA release. (c) Comparisons of the Mfold-predicted structure and the alternative structure predicted from DMS base-pairing data for CORT1. Bases circled in blue or green are sets of bases for which we see support for base-pairing from our DMS data. (d) Overlay of DMS data onto our alternative predicted CORT1 structure. Bases predicted to be unpaired are coloured by base essentiality derived from single mutation data (red: low SSA activity, white: high SSA activity). Paired bases are coloured by essentiality of that specific base-pair, as derived from double mutation data (blue: base-pair is not interchangeable and changing to any other base-pairing results in low SSA activity, white: base-pair is interchangeable and similar activity to parental is observed after substitution with any other Watson-Crick base pair. The invariant universal stem is coloured in grey as there is no corresponding DMS data for those bases.

**Supplementary Table 1.**
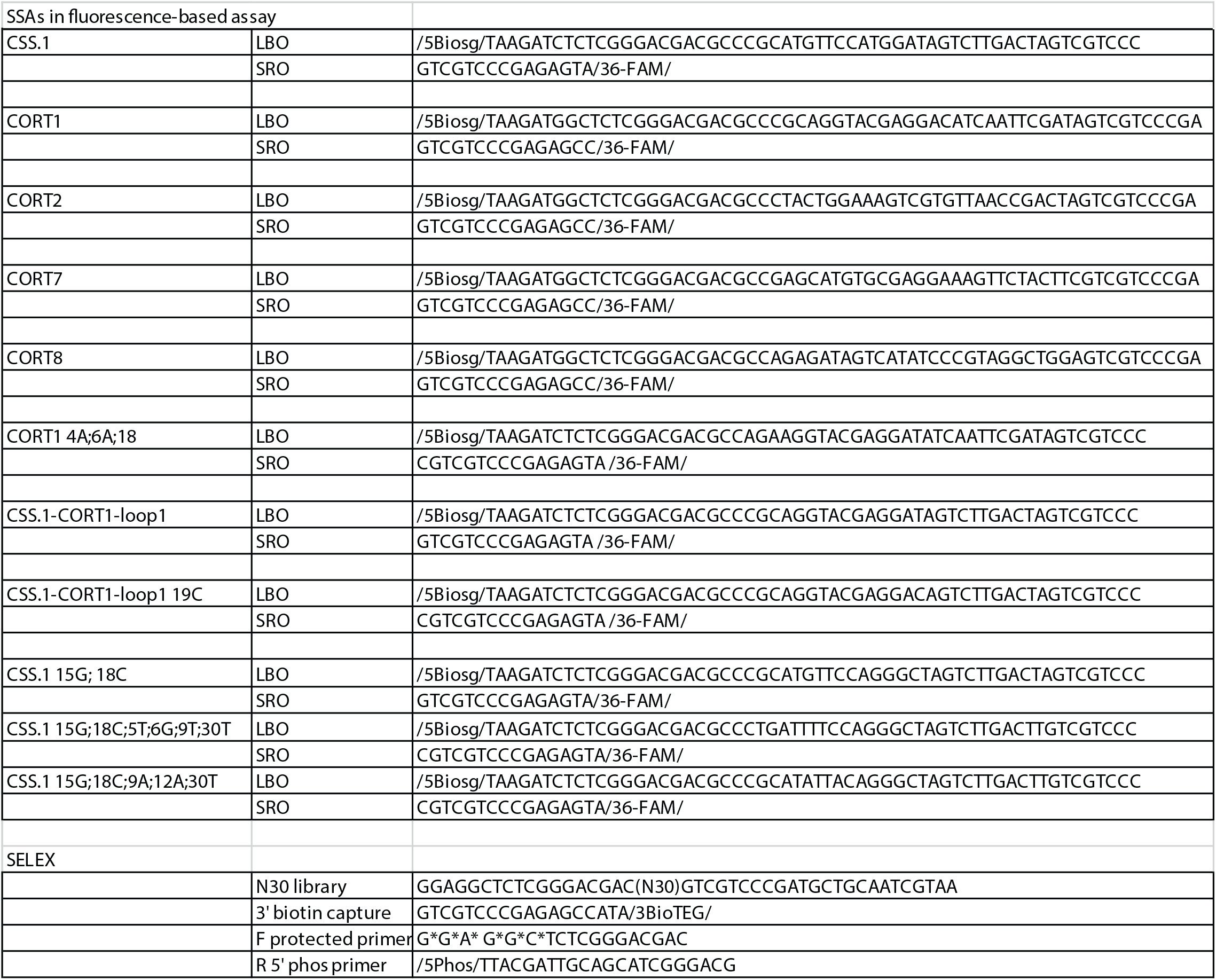
Oligonucleotide sequences of SSAs tested.

